# Identifying and engineering the molecular origins of lignin color for predictive staining of plant tissues

**DOI:** 10.64898/2026.08.03.742431

**Authors:** André Gündel, Delphine Ménard, Marije Nillessen, Antoine Champagne, Leonidas Matsakas, Mika H. Sipponen, Scott E. Sattler, Shinya Kajita, Edouard Pesquet

## Abstract

Lignins in plant biomass are carbon-negative aromatic biopolymers which hold tremendous potential as multipurpose resources for sustainable bioeconomy, limited only by their chemical heterogeneity. Plant lignified tissues, such as sapwood and seed coats, vary in colors within and between species, indicating that specific lignin topochemistries determine the different colors. Yet, the responsible lignin chromogen(s) are unknown. We developed chemical imaging using UV-Vis microspectroscopy to link lignin color to topochemistry in isolates and plant samples. Using synthetic and technical lignins, we identified the different stable chromogens as homomeric lignin substructures varying in size, unit chemistry and interunit linkages. We controlled the accumulation of specific lignin chromogens using genetic engineering to similarly stain lignified tissues from different plant species. We established plant tissue engineering to cast plant tissues with pre-determined color by adjusting lignin topochemistries. We proved that biotechnological manipulation of the identified lignin chromogens predictably and stably stains lignified plant tissues.

## INTRODUCTION

More than 30% of the carbon in the biosphere is stored in lignins^1^, the most abundant renewable source of aromatics^2^ ideal for the biobased economy^3^. However, lignins are heterogeneous polymers which vary differently with cell wall layers, cell types, organs, plant species, and growth/environmental conditions^4^. Lignin heterogeneity is necessary for the physiological properties of each cell wall layer, cell type and tissue. This includes UV-protection for dermal tissues, intercellular cohesion for tissues, mechanical support for wood fibers, pressure-responsive deformation for wood vessels, or intercellular sealant for seed coats^4–5^. In contrast to cellulose made of one unit linked by one linkage, lignins have up to 49 units with 2 to 6 different linkage options^6–7^. Mainly formed with phenylpropanoids, lignins are made non-cell autonomously directly in cell walls by (i) the extracellular release of monomers into cell walls and (ii) their radical oxidation by cell wall embedded PHENOLOXIDASEs: H_2_O_2_-dependent PEROXIDASEs (PRXs) and O_2_-dependent LACCASEs (LACs)^8^. Every monomer shares a common *para* -OH but has distinct aromatic *meta* substitutions (either -H, -OH or -OCH_3_) and different aliphatic chain functions (either -CH_2_OH, -CHO or -COOH)^1–2,4–5^. Lignin polymers extend with the combinatorial formation of covalent bonds between two radicals released by PHENOLOXIDASEs, and the interunit linkage directly depends on the resonating radical forms interacting^1–2,4^. Interunit linkages mainly include (i) the abundant uncondensed aryl-ether linkage between the aromatic *para O* of one unit and the aliphatic chain of another, (ii) the less abundant condensed phenylcoumaran/benzofuran linkage between the aromatic *meta* C of one unit and the aliphatic chain of another or (iii) condensed resinol linkage in-between the aliphatic chains of two units^1–2,4^. The different cell wall layers for each cell type immobilize different PRX and LAC isozyme combinations enabling the accumulation of distinct lignin topo- chemistries differing in (i) accumulation sites, (ii) concentration, (iii) molar weight, (iv) unit chemistries and (v) interunit linkage types^1,4,8^.

The use of lignins as biobased polymeric resources for industry - e.g. in plastic materials^9–11^, in sun-screen lotions^12–17^ or in hair conditioner^18^ – is limited by their color. Technical lignins are generally dark brown to black^19–20^, drastically differing from the color of native lignins in wood where black color is rare^21–22^. Delignification using alcoholysis followed by bleaching^23–24^ completely whitens both hard and softwoods through ring-opening oxidation and fragmentation of aliphatic chains. Lignins thus include multiple chromogens responsible for staining wood differently between species. The dark colors of technical lignins combine native chromogens with others formed during isolation. Non-native chromogens derive from aldehyde-baring substructures formed during mechanical pulping, stilbenes from condensed linkages during alkaline/soda and kraft pulping, chalcones during kraft pulping and various catechols/quinones formed in all isolation processes^19–20^. Reduction of molar weight (M_w_) distribution below 1200 Da “whitens” technical lignins^13,17^, indicating that lignin chromogens are associated with higher M_w_. Yet the molecular substructures determining these different colors in lignins remain unknown. Here, we (i) identified the chromogenic substructures in native lignins, and (ii) showed that their manipulation predictably stained plant tissues. To do so, we established a new chemical imaging method to link color to topo-chemistry using the absorbance of non-solubilized lignins isolated or directly in plant samples using transmission UV-Vis micro- spectroscopy. The identified chromogenic substructures were manipulated by (i) using genetic engineering to increase specific chromogens and predictably change the color of lignified tissues in several plant species and (ii) using newly-developed tissue engineering to assemble plant tissues stained on-demand with specific lignins. The stable color of lignins isolated or incorporated in plants depended on homomeric substructures with distinct extinction coefficients, differing in oligomeric size, unit chemistry and linkage type, which can be controlled to predictively stain plant tissues.

## RESULTS

### Method for measuring biopolymer color

The color of natural biopolymers determines both their physiological roles and their industrial uses. Biopolymer color depends on three factors^25^: (i) its chromogenic transmissive properties linking chemical structure to visible light (Vis) transmittance/absorbance, (ii) its light reflective/scattering and interference properties associated to material surface properties, and (iii) the illumination source. To represent color changes, we converted Vis transmittance/absorbance into RGB (red/green/blue) and HSL (hue/saturation/luminescence) color spaces, following the International Commission on Illumination criteria. To physically measure lignin colors, we developed a new method combining (i) a diamond windowed compression chamber to improve transmittance and homogenize surface properties with (b) transmission UV-Vis micro-spectrometer with variable apertures to allow measurements through solid particles and avoid irregular surface/side effects and (c) illumination with deuterium/halogen lights (Figure S1). This set-up allows (i) measuring the transmission absorbance spectra from UV to NIR of any insolubilized samples (hydrated or not), (ii) eliminating surface reflectance/scattering/interference, and (iii) standardizing color calculation to specific light source. The absolute colors of biopolymers are measured in a few seconds from their transmittance/absorbance converted into HSL color space with hue (from 0- 360°) providing the pure color, saturation (Sat.) characterizing the color intensity, and luminescence (*L) indicating the color lightness/darkness (Figure S1).

### Unit chemistry alters the color of synthetic lignins

The impact of changing lignin unit chemistry on color was evaluated using biomimetically made synthetic lignin homopolymers: dehydrogenation polymers (DHPs). Monomers were externally supplied to purified PHENOLOXIDASEs^26^, but varied in (a) their ring *meta* groups from unsubstituted *p-* hydroxyphenyl (H), monohydroxylated caffeyl (C), monomethoxylated guaiacyl (G) to dimethoxylated syringyl (S) and in (b) their aliphatic chain function with carboxylic acid (- COOH), aldehyde (-CHO) or alcohol (-CH_2_OH). Spectra of monomers and purified DHPs absorbed in the UV range (200-380 nm), peaking at 294-343 nm, but not in NIR (750-1000 nm)^27^. Vis spectra (380-750 nm) differed with lignin unit chemistry by extending absorbances towards the red wavelength range stably with storage time, only observed in DHP polymers but not in most monomers (Figure 1 and S2). In polymers, mid-maximal Vis absorbance wavelength varied with unit aromatic chemistry with H-ringed the least extended (λ_AVG_ = 410 nm) compared to G-ringed (λ_AVG_ = 468 nm), S-ringed (λ_AVG_ = 510 nm) or C-ringed (λ_AVG_ = 520 nm) (Figure 1). Polymers Vis mid-maximal absorbance also differed with unit aliphatic functions, the least for -CHO (λ_AVG_ = 445 nm) and -CH_2_OH (λ_AVG_ = 449 nm) compared to - COOH (λ_AVG_ = 530 nm) (Figure 1). Principal component analysis (PCA) of DHP absorbances revealed three principal components (PCs) that could accurately reconstitute the observed stable colors (Figure S3). PCA grouped the DHPs from light to dark and from white to red, passing through yellow and orange, due to the combined effect of unit aliphatic and aromatic chemistries (Figure 2). Hierarchical clustering of absorbances and HSL values of DHPs formed four clusters with distinct hues from white/grey, yellow, orange to red varying in both brightness and luminescence (Figure 2). Each hue regrouped multiple chromogens with distinct chemistries: S-rings and -COOH for red, G-rings and -CHO for yellow, -COOH functions for orange, and H-rings and -CH_2_OH for white/grey (Figure 2). Overall, lignin chromogens are homooligomeric substructures with different hues determined by unit chemistries.

**Figure 1.**
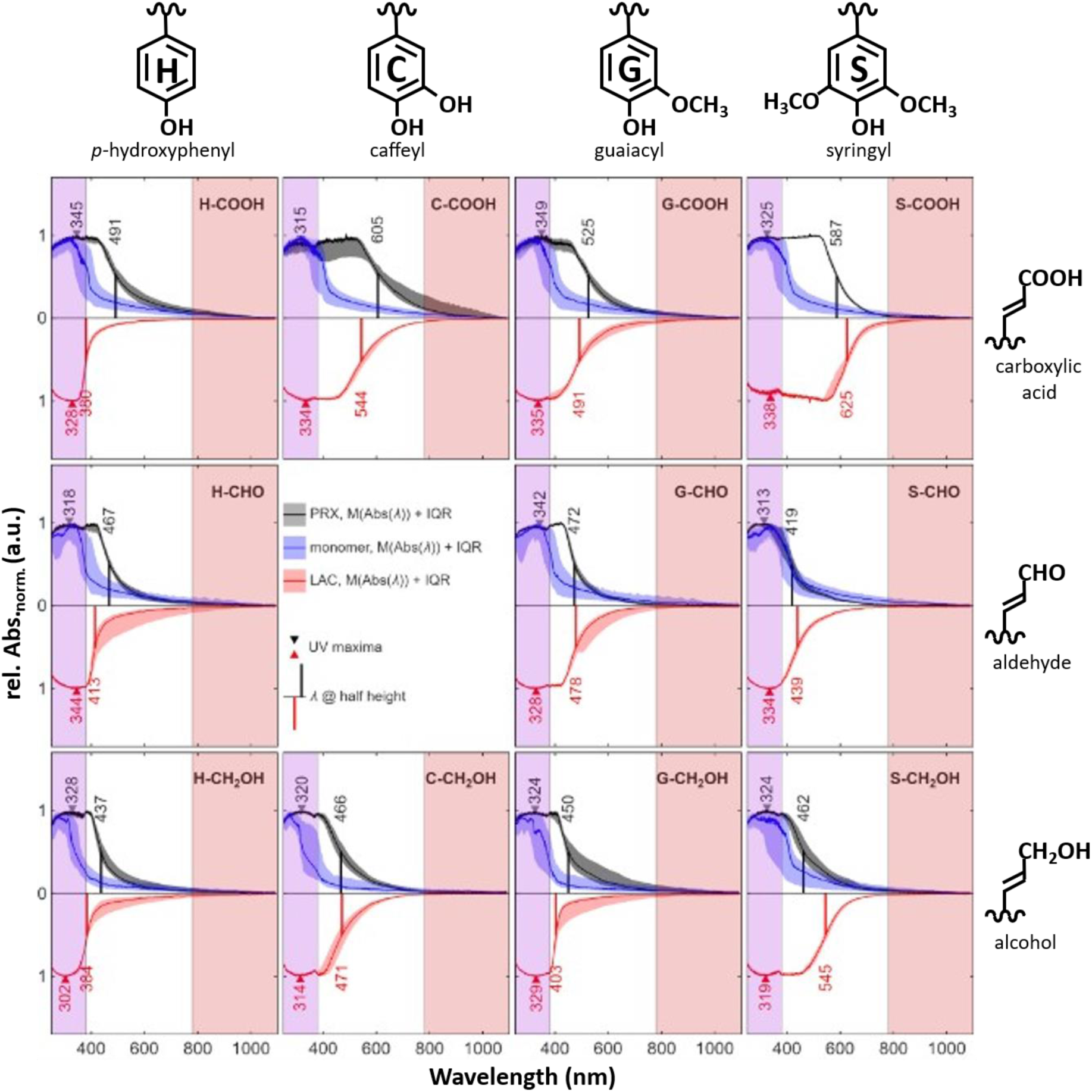
Unit aliphatic and aromatic chemistries combined with specific interunit linkage types change the chromogenic properties of synthetic lignin polymers as analyzed using UV- Vis microspectroscopy. *In vitro* synthetic lignins, called dehydrogenation polymers (DHPs), were biomimetically made by gradually delivering phenylpropanoids to isolated PHENOLOXIDASEs, either H_2_O_2_- dependent PEROXIDASE (PRX, black lines) or O_2_-dependent LACCASE (LAC, red lines). Spectra of 30-50 particles were measured from 200 to 1000 nm to show UV (shaded in purple), Vis (shaded in white) and NIR (shaded in red) absorbances. Synthetic lignins topochemisties were changed from unpolymerized monomers (blue lines) using either different enzymes (PRX or LAC) and chemically distinct monomers varying either in *meta* substitution of aromatic rings (including non-substituted *p-*hydroxyphenyl H, monohydroxylated caffeyl C, monomethoxylated guaiacyl G or dimethoxylated syringyl S) and in aliphatic terminal function (including carboxylic acid -COOH, aldehyde -CHO or alcohol -CH_2_OH) as indicated on the top and left-hand sides of the figure. Panels show the median of range-normalized absorbance spectra of DHP particles and shaded areas represent their interquartile range (IQR). The extent of the red-extended emission is indicated by the wavelength at half height indicating the perceived color. The UV maxima are indicated by triangles, calculated from the weighted average of the upper 1% of absorbance. Vertical lines indicate the mid-maximal Vis absorbances in the electromagnetic spectrum. The presented average spectra for DHPs, monomers and dimers are available in Table S1.

**Figure 2.**
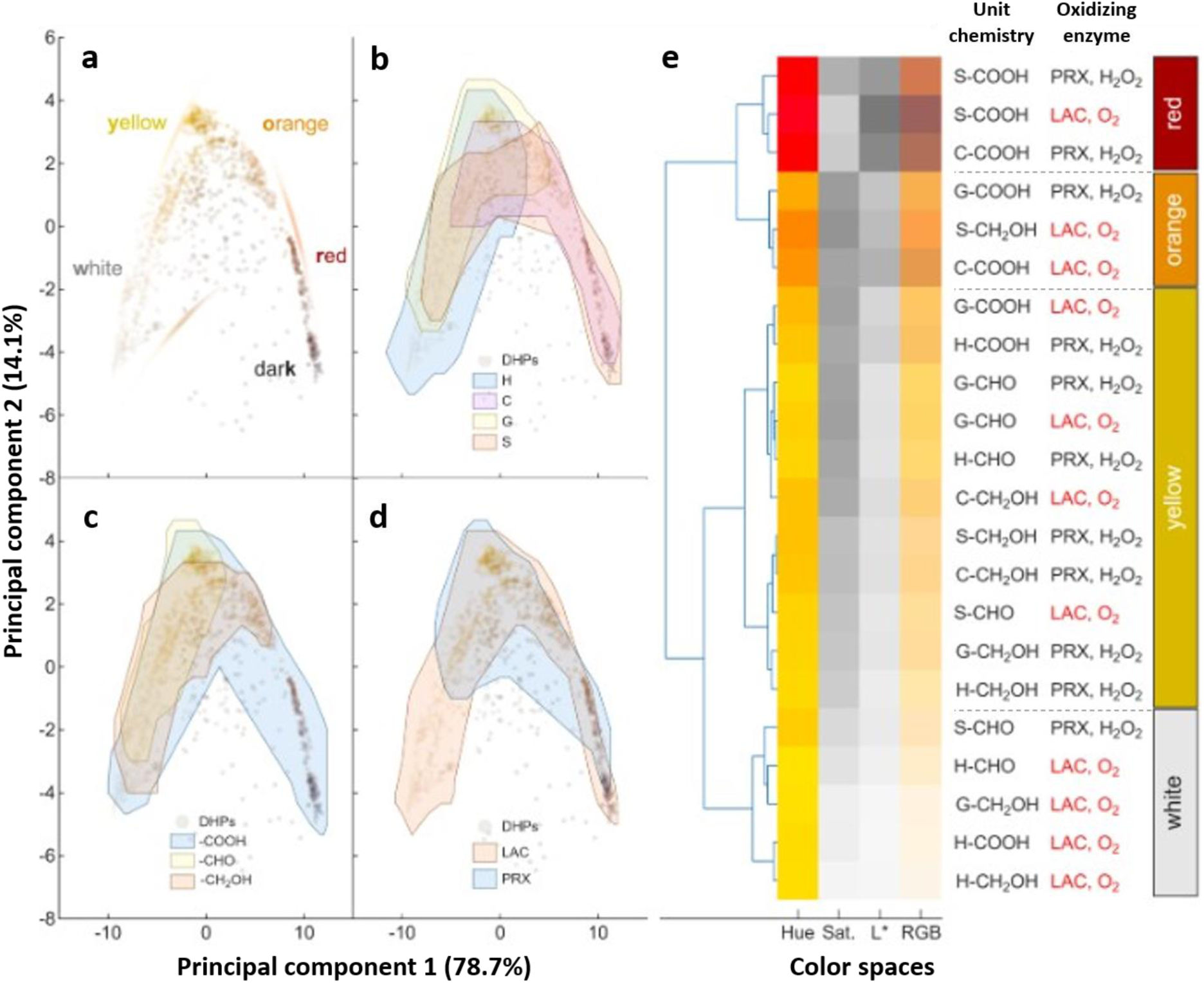
Lignin topochemistries (variations in lignin unit chemistry and interlinkage types) predictively control the colors of these phenolic biopolymers. (a-d) Principal component analysis (PCA) of the absorbances of synthetic lignin polymers onto the two most explanatory principal components (PCs) showing color differences (a) and regions regrouping 50% of synthetic polymer chemistries based on common specific aromatic chemistries (b), aliphatic chemistries (c) and enzyme used (d). Note that different lignin topochemistries, due to unit chemistry and oxidative assembly, are associated to specific colors. Relative contributions and reconstructions of colors from the identified principal components (PCs) are available in Figure S3. (e) Hierarchical clustering analysis (HCA) of synthetic lignin polymer absorbances and corresponding conversions into the hue, saturation (Sat.) and luminescence (L*) and RGB (red/green/blue) color space. Synthetic lignin polymers are grouped into four distinct chromogenic clusters including **white** (grouping 60% of H-ringed units, 80% oxidized by LAC, similar -CH_2_OH and -CHO), **yellow** (grouping similar H-/C-/G-/S-rings, similar -CH_2_OH and -CHO, 60% oxidized by PRX), **orange** (grouping similar C-/G-/S-rings. 65% of -COOH and oxidized by LAC) and dark **red** (grouping 65% of S-ringed unit, -COOH and oxidized by PRX). Differences in absorbances converted in the HSL color space indicated that the color of lignin polymers depended on variations of hue (from yellow to red), of saturation (presented as grey scales corresponding from light to black to color intensity) and of luminescence (presented as grey scales corresponding from light to black to color darkness). Conversion of absorbances to the RGB color space indicated the color of the different lignin synthetic polymers.

### Interunit linkage type alters the color of synthetic lignins

The impact of changing interunit linkages on lignin color was made with DHPs by modifying linkage proportions using different PHENOLOXIDASEs, as shown for G-CH_2_OH forming more condensed linkages with LAC or more uncondensed linkages with PRX^28–29^. DHPs mid-maximal Vis absorbance changed with enzyme-types for each unit chemistry, blue shifted for H and G-ringed units (Δλ_AVG_ = 59 nm) and red shifted for C and S-ringed units (Δλ_AVG_ = 48 nm) independently of aliphatic functions, except for G-CHO and C-CH_2_OH which had the same absorbances with LAC and PRX (Figure 1). The similarity between DHPs made of C-CH_2_OH units reflected the benzodioxane linkage preferentially formed between these units^30^. Condensed and uncondensed G-ringed dimeric model compounds showed no differences in Vis absorbances (Figure S4), indicating that lignin chromogens are larger than the tested model dimers. Mid-maximal absorbance differences between DHPs made with the same unit but by distinct enzymes confirmed differences in interunit linkage proportions. PCA and hierarchical clustering of these DHPs showed that interunit linkage types due to PHENOLOXIDASEs altered the lignin hue with red including DHPs oxidized by as many LACs and PRXs, yellow with higher proportions of PRXs, orange and white/grey with higher proportions of LACs (Figure 2). Altogether, lignin chromogens are homooligomeric substructures with hues determined by interunit linkages.

### Lignin chromogens are oligomers

The impact of oligomer size on lignin color was evaluated by measuring absorbances of DHPs at each degree of polymerization (DP) with size exclusion chromatography. Calibration of retention time (Rt) to molar weight enabled determining the Rt for each DP increment using the hydrodynamic volume of isolated monomers. Vis/UV absorbance ratio between DHPs at each DP increment showed differences in their red-extended absorbance. Chromogenic shifts plateaued at different DP depending on DHP unit chemistry from dimers for G-CHO, to hexamers for S-COOH and to heptamers for H-CH_2_OH (Figure 3). Chromogenic shifts also depended on the oxidizing enzyme used: for similar DP, low variations were observed between -CHO/-CH_2_OH polymers and greater between -COOH polymers (Figure 3). The different absorbance ratios indicated changes in extinction coefficients for each color, differing in saturation such as between S-CHO and G-CHO dimers (Figure 3). Overall, lignin stable chromogens are distinct substructures differing in extinction coefficients due to specific unit chemistry, interunit linkages and oligomeric size. The established method and color palette of lignin hues thus enables determining the lignin topochemistry, homogeneity and color between particles/sites of most samples.

**Figure 3.**
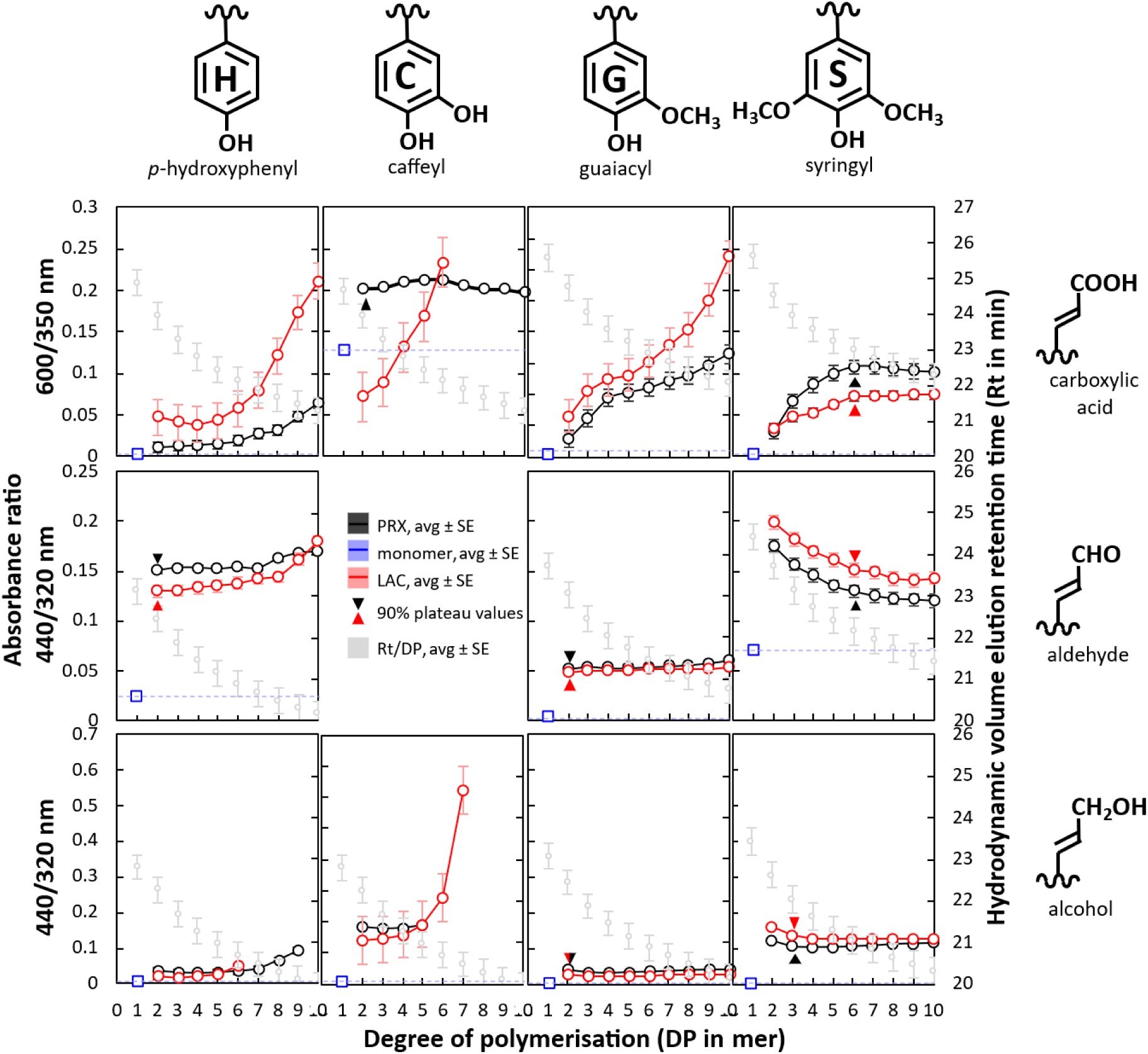
Size exclusion chromatography coupled to UV-Vis spectrophotometry shows that the lignin chromogens vary in oligomeric size and in extinction coefficient with unit chemistry and interlinkage types. Average (avg) size elution profiles of 3-5 technical replicates of 2-3 independently synthesized DHPs solubilized in dimethylformamide (DMF) + 1% LiCl along increasing degree of polymerization according to the measured hydrodynamic volumes of unpolymerized monomers. Retention time (Rt) of isolated monomers and their DP increment from the calibration curve are indicated in grey for each unit chemistry. Error bars indicated standard error (SE) between the different replicated analyses. Example of elution profile monitoring by absorbance spectra is presented in Figure S5. Absorbance ratio between Vis and UV wavelengths, taken from UV-Vis microspectrometry (Figure 1), were used to determine the oligomeric size of the chromogens, relative extinction coefficient (strongest close to 1) and color saturation when reaching 90% of the plateau values as indicated by triangles. Note that some polymers reach 90% plateau at different oligomeric sizes whereas others have their chromogenic capacity increasing with DP. Note that absorbances are affected by solubilization state and solvents making the spectrophometric properties of the same DHPs different between insoluble particles and solubilized in DMF, thus preventing direct transposition of the relative difference in extinction coefficients to lignin isolates and hydrated in plant samples.

### Native lignin colors can be predictively changed using genetic engineering

To demonstrate that the identified chromogens controlled the stable color of plant lignified tissues, we used genetic engineering to specifically increase the proportion of G-CHO dimers conferring a yellow hue in DHPs independently of the enzyme. Redirecting the lignin unit metabolism in *Arabidopsis thaliana* wild-type (WT) plants was made using loss-of-function mutants in (i) *FERULIC ACID-5-HYDROXYLASE-1* (*FAH1*) to remove S-ringed units, (ii) *CINNAMYL ALCOHOL DEHYDROGENASE 4* (*CAD4*) and *5* (*CAD5*) to increase -CHO instead of - CH_2_OH, and (iii) *CAD4 CAD5 FAH1* to remove S-ringed units and increase -CHO. Biochemical analyses confirmed the absence of differences in total lignin content but changes in lignin unit chemistry, with no S-ringed units in *fah1* and large increases of -CHO in *cad4 cad5* (Figure S6)^31–32^. Native lignins were isolated enzymatically from the cell wall polysaccharides in the different genotypes and their absorbances confirmed that lignin color depended on unit chemistry, with both *cad4 cad5* and *cad4 cad5 fah1* colored in yellow similarly to DHPs made of G-/S-CHO whereas WT and *fah1* were white/grey similarly to DHPs made of G-/S-CH_2_OH (Figure 4 and S7). Depletion of S-ringed units with *fah1* did not change the absorbance spectra when stacked with *cad4 cad5* (Figure 4), showing differences in extinction coefficient and/or abundance of S-CHO dimers compared to G-CHO (Figure 3). Native lignins isolated enzymatically have stable colors, due to the same chromogens identified in DHPs, which can be predictively controlled using genetic engineering.

**Figure 4.**
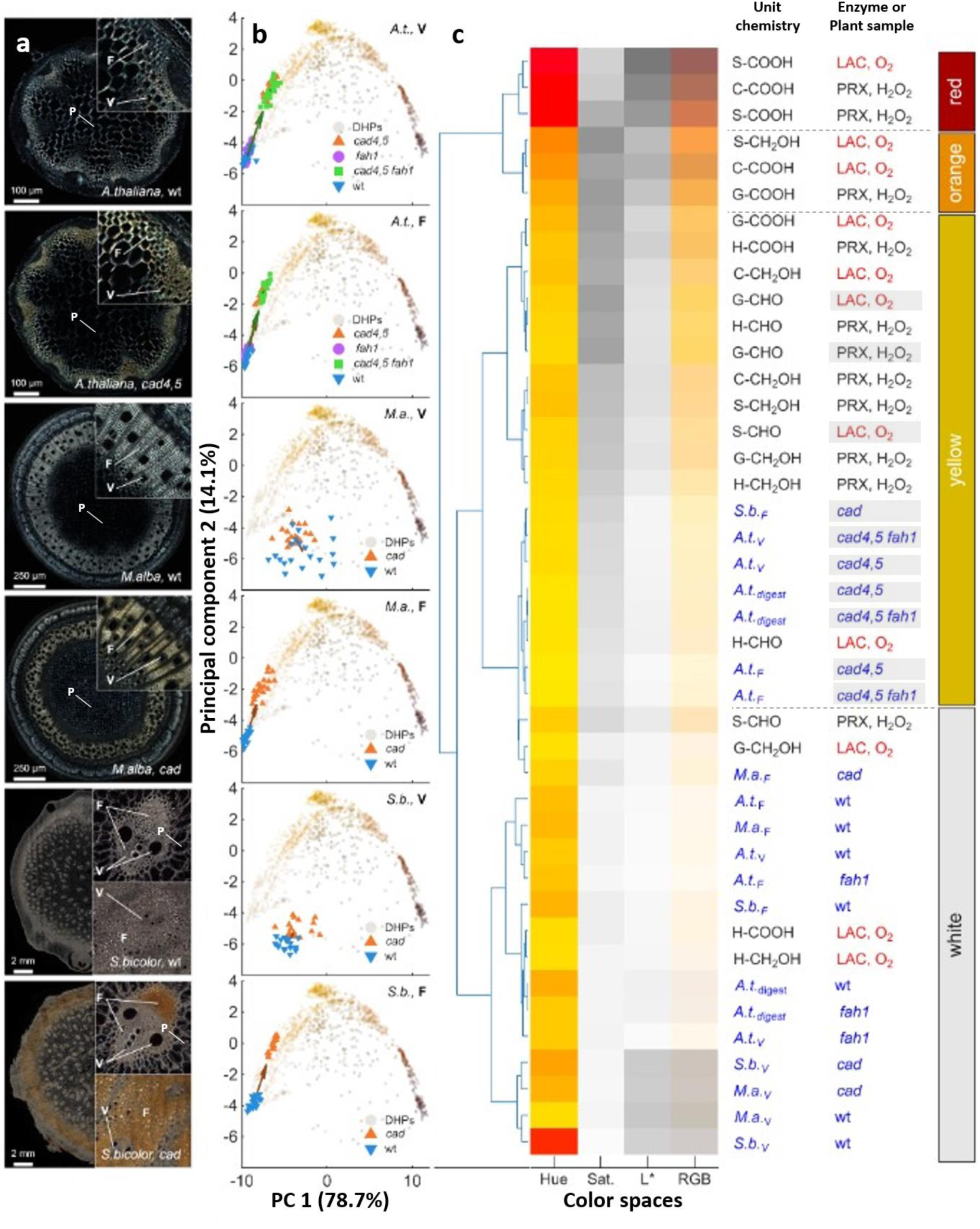
Loss-of-functions in *CINNAMYL ALCOHOL DEHYDROGENASE* (*CAD*) to increase the proportion of chromogenic dimers made of G-/S-ringed -CHO units similarly changes lignified xylem/wood tissue color from white to yellow in different species. (a) Differential interference contrast (DIC) images of cleared stem cross-section between wild- type (wt) and *CINNAMYL ALCOHL DEHYDROGENASE* loss-of-function mutant (*cad*) plants in genetic model thale cress (*A. thaliana*^31–32^), hardwood tree mulberry (*M. alba*^34^) and agricultural crop sweet sorghum (*S. bicolor*^33^), which have respectively speciated apart from each other 120 and 160 million years ago (according to geological times). Images of cross- sections and close-ups of xylem/wood tissues are shown in the upper right corner with lignified fibers (F) and vessels (V) indicated. Note that in contrast to xylem/wood tissues stained in yellow in *cad* mutants and white in wt plants independently of the species, parenchyma cells (P) in piths are similarly colored in white for all genotypes and species. Bars indicate scales for each cross-section. (b) Principal component analysis (PCA) of the absorbances of 20-30 cells compared to the absorbances of synthetic DHP homopolymers for each cell type (vessels V and fibers F) and plant species in wild-types (wt) and different loss-of-function mutants including *fah1*, *cad4 cad5* and *cad4 cad5 fah1* for thale cress (*A. t*), natural *cad* mutant Sekizaisou for mulberry (*M. a*) and EMS brown midrib *cad* mutant *bmr6* for sorghum (*S. b*). Note that both cell types display a red-extended shift towards the yellow color, differing in extent and homogeneity with cell types and species. (c) Hierarchical clustering analysis (HCA) of the absorbances measured *in situ* directly on cell types in organ biopsies for vessel (V) and fiber (F) cells in the different plant species and genotypes (GT) together with enzymatically isolated native lignins from *Arabidopsis* (Figure S7) and the synthetic DHP homomeric polymers. Differences in absorbances converted in the HSL and RGB color spaces showed that the color changes from white to yellow in both cell types and enzymatically isolated lignins in *cad* mutants clustered with synthetic lignins made of G-/S-CHO units (shaded in grey).

### Changing the native color of wood cells and seed coats by redirecting the metabolism

As lignin topochemistry varies between cell types^11,42–43^, we measured absorbances in wood vessel and fiber cells directly in cross-sections (Figure 4). Experimental conditions were optimized using signal-to-noise ratio and working range which showed best performance with compressed samples: reducing reflective/interfering surfaces, improving transmittance, but lowering sub- cellular resolution (Figure S8). To prove that the same chromogens similarly colored lignified tissues in different plant species, we compared WT and *cad* mutants to increase yellowish G-/S-CHO dimers in *Arabidopsis*^31–32^, the cereal *Sorghum bicolor*^33^ and the hardwood *Morus alba*^34^. In all species, cell type absorbances were red-extended in *cad*, changing from white to yellow/orange as observed in xylem of stem cross-sections compared to unchanged whitish in pith parenchyma (Figure 4). Variability depended on cell type and species, more stable in *Arabidopsis* than in other species (Figure 4). Lignins in sorghum samples were removed oxidatively in acidic conditions^35–36^, which completely removed the *cad*-dependent yellow hue in all cell types to white, further confirming lignin color-dependency (Figure S9). Overall, the same genetic engineering strategy can be used to steer the accumulation of specific chromogenic substructures to predictably stably stain homologous lignified cell types/tissues independently of the plant species.

We then measured the absorbances of *Arabidopsis* lignified seed coats^1,4–5^, characterized by a Vis mid-maximal absorbance of 444 nm compared to 371-387 nm for wood cells, reflecting the whitish color of wood compared to the yellowish color of seed coats (Figure S10). We used gain-of-function of *FAH1* to increase S-ringed units^37^ and loss-of-function in *LAC*s^38^ to prove the intervention of different PHENOLOXIDASEs to form distinct chromogenic substructures. Increasing S-ringed units enabled significant color change in one of the two overexpressing lines tested, known to differ in their overexpression levels^37^. Modifying multiple LACs controlling seed coat lignin accumulation^39–40^ significantly shifted seed coat color to more yellowish hues (Figure S10). The different colors of lignified tissues can be genetically controlled by the accumulation of chromogenic substructures made from specific unit chemistry and oxidizing enzymes.

### Technical isolation of lignins alters their colors

We then evaluated the stable absorbances of technical lignins. Both the pulping process and the plant species affected the absorbances of isolated lignins: soda lignins from mixed annual plants were orange, kraft/sulfate lignins from softwoods were yellow, whereas alkaline and sulfite lignins from mixed softwoods were white/grey (Figure 5). PCA and hierarchical clustering of technical lignins with DHPs showed that the color/absorbance corresponded for softwoods to yellow/orange due to their H- and G- ringed units and extending colors to red for grasses due to their S-ring units (Figure 5). Our method showed that among softwoods, the red-extension between kraft, soda and sulfite pulping depended on reduction of -CH_2_OH and an increase of ring methoxylation during pulping (Figure 5 and Table S2). Heterogeneity between particles was greater when using mixed biomass than one species. To prove that both biomass type and chemical treatment affected lignin colors, we compared lignins isolated from beech or spruce sawdust using the same organosolv process with or without 1% sulfuric acid. Absorbance measurements showed that organosolv lignins isolated without acid were white/grey for hardwood and yellow/orange for softwood, whereas acid treatment colored in yellow/orange only hardwoods (Figure 5). Heterogeneity between particles increased following acid addition for both feedstocks (Figure 5). Biochemical analyses showed a large reduction in ring methoxylation levels following acid organosolv only in hardwoods, suggesting acid-induced modification of S-ringed units, including partial demethylation and possible formation of catechols/quinones, potentially contributing to the observed color change. Overall, the lignin isolation procedure, by introducing controllable non-native chemical modifications, offers an additional mean of tuning the color and heterogeneity of lignin particles.

**Figure 5.**
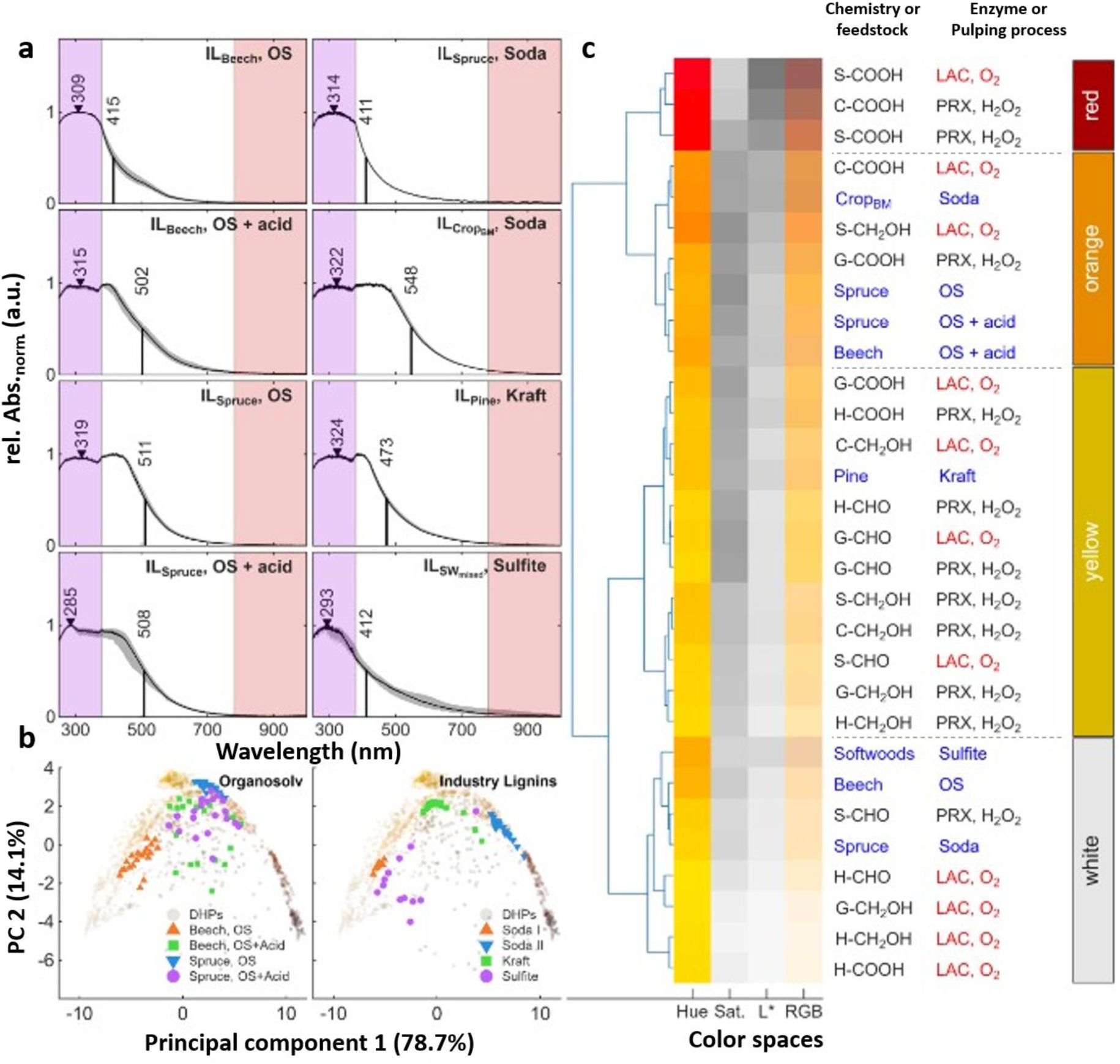
The color of technical lignins depends on both the plant feedstock and the pulping processes, combining native and non-native chromogens, both adjustable to control the final coloring. (a) Average UV-Vis absorbance spectra of 20-30 particles and IQD (in grey shades) of technical lignins made from different feedstocks (single or mixed species) and different processes including organosolv (OS), sulfate/Kraft, sulfite/lignosulfonate or soda pulping. Maximal absorbances are indicated by triangles and mid-maximal Vis absorbance wavelength with black lines. Note that the large effect of both feedstocks (annuals vs hardwoods vs softwoods) and pulping process on the red-extension of the mid-maximal Vis absorbances. (b) Principal component analysis (PCA) of the absorbance spectra of the different technical lignins compared to the DHP lignin color palette showing both the color of the different lignins as well as the variability between their isolated particles. (c) Hierarchical clustering analysis (HCA) of the average absorbance spectra of the different technical lignins compared to the DHP lignin color palette. Differences in absorbances converted in the HSL and RGB color spaces showed the significant effect of the pulping process on the color of softwood (pine or spruce) lignins from white with sulfite/lignosulfonate to yellow with sulfate/Kraft to orange with organosolv.

### Engineered plant tissues predictively colored with specific lignin chromogens

Specific lignin chemistries are necessary for plant physiology, as deviations can otherwise jeopardize growth^1–2,4–5^. The full lignin color palette cannot therefore be controlled in whole plants as easily as for DHPs. We thus developed plant tissue engineering. This is one of the first reports of assembling plant tissues from isolated cells joined together by their cell walls with lignins, similarly to whole plants. To simultaneously steer the accumulation of specific lignin chromogens and join the cells together into plant tissues stained on-demand, we cast wood or ground plant tissues into a mold as agglomerated material combining (a) differentiated plant cell types made using inducible pluripotent cell suspension cultures (iPSCs) triggered with different hormones^41–42^ (Figure S11) with (b) specific monomers^4,5^. Wood and ground tissue-engineered samples with predefined color were cast with either wood vessels or parenchyma cells, each expressing different PHENOLOXIDASEs changing their non-cell autonomous oxidation capacity^43–45^ (Figure 6), specific lignin monomers (H-COOH, G-CH_2_OH, G-CHO or S-COOH), PHENOLOXIDASE inhibitor NaN_3_ and/or H_2_O_2_. Tissue coloring varied with (a) NaN_3_/H_2_O_2_ treatment confirming that each cell types have different LAC/PRX capacity, and (b) monomer chemistry which combinedly controlled the formation mainly in wood tissues of shades of red chromogens with S-COOH, yellow/orange with G-CHO and white with H-COOH like in DHPs (Figure 6). The stable color resulting from engineering lignified plant tissues could thus be predictively steered by controlling cell types, supplied lignin unit chemistry and oxidative enzyme capacities.

**Figure 6.**
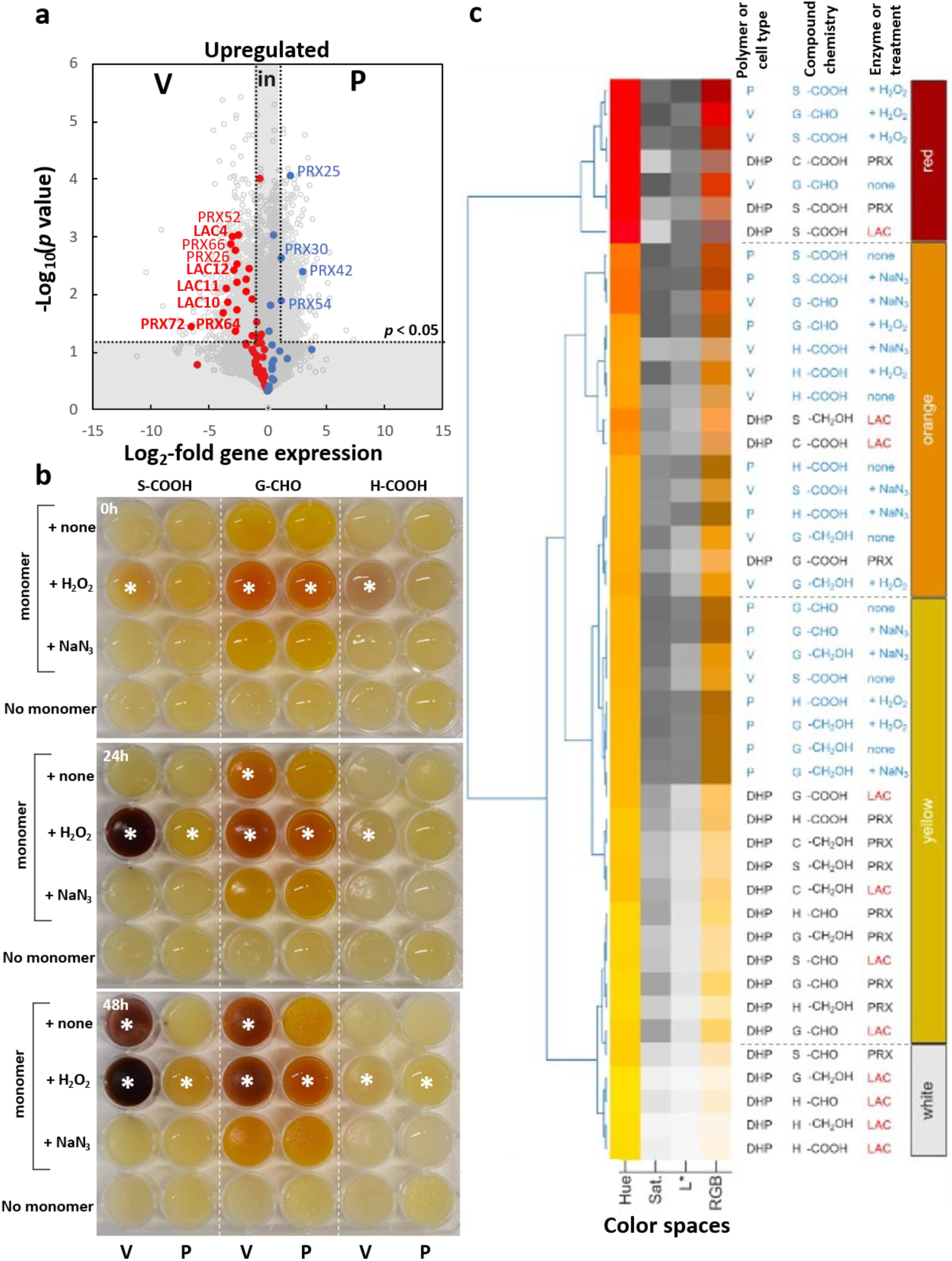
Predictive stable coloring of plant tissues by directing the accumulation of specific lignin chromogens and the cell types using plant tissue engineering. (a) Volcano plot showing differential gene expressions between vessel (V) and parenchyma (P) cell types formed on -demand using inducible pluripotent cell suspensions cultures (iPSCs) and revealing differential upregulation of specific LAC and PRX paralogs in vessel compared to only PRX in parenchyma (significant threshold at absolute log_2_ fold change > 1 and significant adjusted Student T-test, α = 0.05). PHENOLOXIDASEs previously functionally associated with lignin oxidative formation are indicated in bold^1,8,38–40^. (b) Time-course analysis of engineered tissues using iPSCs differentiated either with vessels (V) or parenchyma (P), as described in Figure S11, externally supplied either with no monomer or with monomer together with either H_2_O_2_ or NaN_3_ for S-COOH, G-CHO and H-COOH. An example of cm-wide larger casting into traditional Swedish *Dalahäst* (Dalecarlian horse) made from vessel cells bound together with S-COOH and H_2_O_2_ is presented in Figure S12. Note that oxidative capacity is strictly associated with specific cell types induced in iPSCs, as PRX and LAC-expressing vessels can actively oxidize S-COOH in red and G-CHO in yellow/orange with monomer alone whereas PRX-expressing parenchyma can only oxidize monomers in the presence of H_2_O_2_. Stars indicate significant differences between samples with no monomer or samples with monomer and NaN_3_, both defining cell and unpolymerized monomer background controls (appearing mostly yellow/white). (c) Hierarchical clustering analysis (HCA) of the HSL and RGB color spaces measured for wood and ground engineered tissues (indicated in blue) assembled with different lignin monomers comparatively to synthetic DHP homopolymers (indicated in black). Note that hue, in contrast to saturation and luminescence, have been predictively reached with S-COOH to red and with G-CHO to orange, considering that tissue cell and unpolymerized monomer background controls are colored yellow.

## DISCUSSION

We herein unambiguously identified the chromogens responsible for native stable lignin colors: homomeric substructures differing in extinction coefficient due to specific unit chemistry, interunit linkages and oligomeric size. To reliably measure the absolute color of insoluble lignins in isolates and biopsies, we developed a new chemical imaging method combining diamond windowed pressure chambers with transmission absorbance microspectroscopy, to quickly measure absorbances with <µm resolution more accurately than previously using RGB^46–47^. Our new method follows green chemistry principle by using no catalyst nor harmful solvent. Our method provides crucial information for rapid fingerprinting on the composition and heterogeneity of whole lignins due to red-extended absorbances only observed for polymers and directly reflecting aromatic, aliphatic and interlinkage types. Extending other chemical imaging methods, like atom-resolving X-ray^32^ or chemical group-resolving Raman^31^ microspectroscopies, UV-Vis microspectroscopy detects oligomeric substructures. We anticipate that the correlated use of multiple label-free microspectroscopic approaches will reach unprecedented *in situ* lignin characterization extending the current limitation of chemical imaging^5^.

The chromogens determining the stable color of lignified tissues have been questioned since the mid-1990s, with the first transgenic plants engineered with antisense against *CADs*^48–50^. In contrast to their transient wine-red color due to temporally unstable sinapaldehydes^51^, we herein demonstrate that the stable yellow coloring depends on dimers of G-CHO independently of the plant species. Our genetic engineering strategy proved that the combined manipulation of unit chemistry and PHENOLOXIDASEs enabled us to accurately steer the staining of lignified tissues. Extending the color palette will depend on oxidizing additional phenolic compounds, like anthocyanins for purple wood in *Arabidopsis*^52^ or alkaloids for black in *Diospyros*^20–21^. As manipulation possibilities are limiting tissue/plant physiology/growth, we developed plant tissue engineering using plant tissue cohesion principles (Figure 6). We combinedly used iPSCs to control cell type differentiation and their oxidative capacities, and bound cells into tissues using cell walls’ autonomous capacity, similarly to whole plant tissues^4–5^. Plant tissue engineering unambiguously proved the contribution of specific lignin chromogens adjusting the stable colors of lignified tissues, and represents a methodological breakthrough to understand plant tissue properties. Plant tissue engineering controlling tissue types with predictable adjustable coloring represents promising biotechnology applications for the restoration of historical wood in artefacts and musical instruments.

## METHODS

### Plant material – whole plants and inducible pluripotent cell suspension cultures (iPSCs)

*Arabidopsis thaliana* plants were grown in 4:1 peat soil:vermiculite under long-day conditions: 16:8 light:dark cycles at 22:18°C and 60% relative humidity in growth chambers of Stockholm University (lat. 59°36’ N, long. 18°06’ W). *Arabidopsis* loss-of-function mutants in *CAD4* (SAIL_1265_A06^31^), *CAD5* (SAIL_776_B06^31^), *FAH1* (*ref3-2* EMS mutant^31^) and *LAC4 LAC5 LAC10 LAC12 LAC17*^38^ in the Col-0 accession background were all genotyped to ensure homozygosity of the mutations using PCR as previously described^31–32^. *Morus alba* included natural *cad* mutant “赤材桑/Sekizaisou” and wild-type F1 hybrids from crossing “赤材桑/Sekizaisou” with “国桑21号/Kokusou 21” as previously described^45^. Samples for both genotypes were taken from 2-year-old plants cultivated in pots, outside for “Sekizaisou” and in greenhouse for the F1 hybrid at the Koganei campus of Tokyo University of Agriculture and Technology (lat. 35°70’ N, long. 139°52’ W). All sections were prepared from growing branches at the 6th internode. *Sorghum bicolor* wild-type (B Tx623) and *bmr6*-*ref* B Tx623, which contains a diethyl sulfate (DES) induced nonsense mutation in sorghum *CAD* gene (Sobic.004G071000), were selected as previously described^33,53^. Sorghum plants were grown in the greenhouse on the East Campus of the University of Nebraska – Lincoln during (lat. 40°50’N, long. 96°39’W) winter of 2020-2021. The potting mix was comprised of 38.5% peat, 23.1% soil, 19.2% sand, and 19.2% vermiculite. The greenhouse conditions were set at temperature 30/27 ± 1°C, and light/dark 12/12h. Gain-of-function mutants transformed with *C4H:FAH1* were selected based on segregation of descendants exhibiting antibiotic resistance as previously described^37^. Stem samples for *Arabidopsis*, *Morus* and *Sorghum* were stored in 70% ethanol:water, renewed 3-4 times, at 4°C for 2 months to 5 years until characterization to evaluate the stable color of plant tissues.

Inducible pluripotent cell cultures (iPSCs) from *Arabidopsis thaliana* were used as previously described^41–44^ to induce on-demand the synchronized differentiation of wood cells by adding phytohormones or maintaining actively dividing parenchyma cells without adding phytohormones. Phytohormone treatment consisted of 6 µg/mL α-naphthaleneacetic acid (N0640, Sigma Aldrich), 1 µg/mL 6-benzyl-aminopurine (B3408, Sigma Aldrich) and 4 µM 24-epibrassinolide (E1641, Sigma Aldrich) in fresh full-strength Murashige and Skoog (MS) medium (M0222.0025, Duchefa) at pH 6.0 with 10 µM of morpholino-ethanesulfonate (M8250, Sigma-Aldrich) and 3% (w/v) sucrose supplied to 30 mg mL^−1^ of 10-d-old cell suspension (fresh weight measured after 200 × g centrifugation and removal of supernatant) to synchronize cell cycle to G_0_ due to sucrose depletion during extended growth in plateau phase^41–42^. Cell growth was measured using packed cell volume (PCV) relating pelleted cell volume after 2 min 200 × *g* centrifugation and total volume in 15mL sterile conical tubes, whereas cell differentiation efficiency was monitored after 7 days of culture using differential interference contrast (DIC) using an inverted Zeiss Axiovert 200 microscope equipped with an Axiocam 512c camera and controlled using ZEN 3.1 (ZEN Lite) software (Zeiss, Sweden). Images were obtained using a 10× objective (NA 0.30). Birefringence was measured using the 8-bit grey scale converted color images and the ImageJ software.

### Tissue engineering

Tissue engineering was performed by molding concentrated cell suspensions in multi-welled polystyrene plates or silicon mold. iPSCs, either differentiated into vessel or parenchyma cells to respectively form wood or ground tissues, were centrifuged at 1600 × g for 3 min using a swing-out rotor (A-4-44, Eppendorf) at 23°C in a refrigerated centrifuge (5180R, Eppendorf) and washed 5 times with ultrapure water at 18.2MΩ.cm (Puranity TU+, VWR) in 50mL conical tubes. Final concentrations of cell suspensions were adjusted to 80% PCV in ultrapure water, supplemented with 80µl/mL of 120mM lignin monomers in DMSO together with or without 40µl/mL of 100mM hydrogen peroxide (H_2_O_2_; 95299, Sigma-Aldrich) in water or 50µl/mL of 50mM sodium azide (NaN_3_; S2002, Sigma-Aldrich). Cell suspension slurries were mixed by pipetting using 5mL serological pipettes and then deposited in each well, and left to incubate in a thermoregulated room (23±0.5°C) in the dark. Imaging was performed using a transilluminator (Mini LED Light Box, Fisher Scientific) and a D750 digital camera (Nikon, Sweden) equipped with a 105mm F2.8 EG macro-lens (Sigma, Sweden).

### Chemical reagents, technical lignins and dehydrogenation polymer (DHP) synthesis

Phenolic compounds used included t-cinnamic acid/P-COOH (C80857, Sigma-Aldrich), *p*- coumaroyl alcohol/H-CH_2_OH (PHL82506, Sigma-Aldrich), *p*-coumaroyl aldehyde/H-CHO (C755450, Toronto Research Chemicals), *p*-coumaroyl acid/H-COOH (55823, Sigma-Aldrich), coniferyl alcohol/G-CH_2_OH (223735, Sigma-Aldrich), coniferyl aldehyde/G-CHO (382051, Sigma-Aldrich), ferulic acid/G-COOH (W518301, Sigma-Aldrich), sinapyl alcohol/S-CH_2_OH (404586, Sigma-Aldrich), sinapyl aldehyde/S-CHO (382159, Sigma-Aldrich), sinapic acid/S-COOH (D7927, Sigma-Aldrich), caffeic acid/C-COOH (020M1309, Sigma-Aldrich), caffeyl alcohol/C-CH_2_OH (PHL82489, Sigma-Aldrich). Dimeric models included pinoresinol/β-β_G2_ (40674, Sigma-Aldrich), guaiacylglycerol-β-guaiacyl ether/β-O-4_G1_ (CDS013307, Sigma-Aldrich), dehydrodiisoeugenol /β-5_G2_ (PHL83844, Sigma-Aldrich) and 2- phenyl-benzofuran/β-5_P2_ (S49779, Sigma-Aldrich).

Technical lignins included soda-alkaline lignins from spruce (370959, Sigma-Aldrich), soda- alkaline lignins from mixed annual plant biomass (including sugarcane bagasse) (Protobind 2400, GreenValue Enterprises LLC, Media PA, USA), kraft lignins from softwood (pine) (BioPiva 100, UPM, Finland) and lignosulfonate from mixed softwoods (Lignosulfonate DS10, Domsjö Dabriker AB, Sweden). Organosolv lignins were isolated from beech or spruce sawdust as previously described^54^. Briefly, beech sawdust was treated in a 60% (v/v) ethanol/water solution at 180°C for 60 min, whereas spruce sawdust was treated in a 50% (v/v) ethanol/water solution at 200°C for 30 min. Both feedstocks were processed in the absence or presence of 1% (w/w_biomass_) sulfuric acid at a liquid-to-solid ratio of 10 (mL/g). Following organosolv treatment, the cellulose-rich pulp was separated from the slurry by vacuum filtration. Ethanol was subsequently removed from the liquid fraction by rotary evaporation, after which the lignin was recovered by centrifugation (10,000 × g, 15 min, 4 °C), dried, and stored at room temperature. Biomimetically synthesized lignin DHPs were made *in vitro* according to the *Zutropf* method, as previously described^15^, where 10 ml of a PHENOLOXIDASE solution, either with 10 mg of horseradish PEROXIDASE (P8375, Sigma-Aldrich) or with *Trametes* LACCASE (38429, Sigma-Aldrich) solubilized in 0.1 M NaH_2_PO_4_ (13274, Merck) buffer at pH 6 was mixed under magnetic stirring. Gradual supply at a rate of 0.5 mL h-1 using a syringe pump (model 101, KD scientific) of a 10 mL solution with 12 mM of monomer in 3:7 methanol:0.1 M NaH_2_PO_4_ buffer at pH 6 was made in addition to either 10 mL solutions of 14 mM H_2_O_2_ (95299, Sigma-Aldrich) for PEROXIDASE or bubbling air using an aquarium pump for LACCASE. After 24 h, DHPs in the mixture were precipitated by adding 150 µL 12 M HCl (1.00317, Merck) and incubating 1h at 4°C, then centrifuged at 10,000 × g using a fixed angle rotor (FA-45-6-30) in a refrigerated centrifuge (5180R, Eppendorf) at 4°C for 30 min, the supernatant was removed, and the pellet was washed two times with 1-butanol (281549, Sigma-Aldrich), two times with ultrapure water and finally freeze-dried. DHP produced were stored on the bench in closed microtubes for several months prior to characterization.

### Size exclusion chromatography (SEC)

Monomers and DHPs were solubilized at weight concentration of 3.3 to 5 mg/mL in a solution of dimethylformamide (DMF) with 1% LiCl (L9650, Sigma-Aldrich) and incubated for two days at 50°C under inversion in a mini-labroller (Labnet, USA) placed in an INCU-line IL23 (VWR, Sweden). Solubilised supernatants were transferred by pipetting into glass insert and sealed in glass tube with pierceable caps. Around 13 to 15 equivalent µg of each samples (5 to 3 µL) were analyzed by size exclusion chromatography (SEC) using a Prominence LC system (Shimadzu, Japan) on a PSS GRAM column (10 µm, 8 × 300 mm)/PSS GRAM precolumn (10 µm, 8 × 50 mm) kept at 50°C with a mobile phase made of DMF + 1% LiCl at a gradually reducing flow rate from 0.6 mL/min to 0.3mL/min in 70 min. Elutions were monitored using absorbance spectrophotometry in cells at 50°C with a SPD-M20A diode array detector (Shimadzu, Japan) under halogen and deuterium lamps illumination from 200 to 800 nm (slit width of 1.2nm). Measured retention times (Rt) varied between 15 and 35 min, leaving the additional 35 min to wash the column prior to the next injection. Determination of Mn, Mw and Mz were made using LabSolutions GPC Software v5.87 (Shimadzu, Japan) calibrated using ReadyCal-Kit Poly(styrene) low (PSS-pskitr4l) with 14 distinct polymers with Mw ranging from 266 to 66,000 Da (PSS Polymer Standards Service GmbH, Germany). Exponential elution curve, relating Rt to Mw, was determined using five independent replicates and degree of polymerization (DP) of each DHP was determined from the hydrodynamic volume measured for each isolated monomer.

### Gene expression analysis

Gene expression was performed using data (GSE73146) previously published by the corresponding author on iPSCs differential gene expression (n = 3 independent replicates) between living wood vessels and living parenchyma cells. Volcano plots were performed comparing adjusted Student t-test *p-*values with log_2_ fold gene expression ratio. Significance threshold for differential gene expression was set for |log_2_ fold gene expression ratio| > 1 and *p* value < 0.05. Locus numbers of *Arabidopsis* LACCASE and PEROXIDASE genes were taken from^1,8,38^.

### Plant sectioning, maceration and image acquisition

The basal 2 cm of the stems were stored in 70% (v/v) ethanol at –20°C until sectioning. The stem bases of *Arabidopsis* and mulberry were vacuum-infiltrated with ultrapure water, embedded in 10% (w/v) agarose (A9539, Sigma-Aldrich), and sectioned to 50-µm thickness using a Leica VT1000s vibratome, whereas sorghum stems were embedded in cryo-embedding media (FSC22 Clear, Surgipath), frozen at -20°C and cryo-sectioned at 50-µm thickness using a Leica Cryostat E342.

Oxidative lignin removal in acid conditions of sorghum tissues was performed as previously described^35–36^. In brief, plant materials were transferred to microtube with a safety lock, and supplemented with maceration solution (3.5% H_2_O_2_ in 50% glacial acetic acid (1.01830, Supelco) in ultrapure water). Samples were incubated in a chemical fume hood at 95 °C for 3 hours in a heat-block (drybath, ThermoScientific). After incubation, tubes were vortexed horizontally for 1h (Vortex genie 2 mounted with horizontal tube holder) and then left to sediment for 15 min. Excess maceration solution was removed by pipetting and a solution of 10% Na_2_CO_3_ (71360, Sigma-Aldrich) in water was added to neutralize pH. Cells were further mechanically separated using pipetting.

DIC images of cross-sections and isolated cells were acquired using a Zeiss Axiovert 200 microscope equipped with an Axiocam 512c camera and controlled using ZEN 3.1 (ZEN Lite) software. Cross-sections were imaged using a 5x objective (NA 0.15) and tiled using ImageJ software Fiji^55^. Cell macerates were imaged using a 10x objective (NA 0.30) and analyzed using a custom macro for semi-automated detection of region of interest (ROI) based on intensity thresholding (8-bit conversion and particle analysis) in ImageJ Fiji (Supplementary Data 1). Cell characteristics, including hue, were quantified after conversion to an HSB image and measurements were made within defined ROIs. Seeds were imaged using a VWR VisiScope ZTL350 stereomicroscope equipped with a VISICAM TC 10 camera (WVR). Additional images were captured using a Samsung Galaxy S24 smartphone.

### Cell wall isolation and enzymatic removal of cell wall polysaccharides

Cell wall isolation from stems of the different *Arabidopsis* genotypes for 10-week-old plants was performed as previously described^26^. Once harvested, plant material was rapidly frozen in liquid nitrogen and ground to a fine powder using a ceramic pestle and mortar. Approximately 100 mg of powdered tissue was mixed with extraction solution (140 mM Tris base (Sigma-Aldrich, T1503), 105 mM Tris-acetate (Sigma-Aldrich, T1258), 0.5 mM EDTA (Scharlau Chemie, AC0965) and 8% (w/v) lithium dodecyl sulfate (Sigma-Aldrich, L4632)), vortexed and centrifuged at 10,000 × g for 10 min. This extraction step was repeated twice more, followed by two washes with ultrapure water, two washes with methanol, one wash with chloroform:methanol (1:1, v/v) and three washes with acetone. Pellets were air-dried overnight. Approximately 10 mg of dried cell wall material was incubated in cell wall polysaccharide enzymatic digestion solution containing 2.65 mg/mL of CELLULASE (C9748, Sigma-Aldrich), 0.95 mg/mL of β-GLUCOSIDASE (49290, Sigma-Aldrich), 2.65 mg/mL of HEMICELLULASE (H2125, Sigma-Aldrich), 2.65 mg/mL of XYLANASE (X2763, Sigma-Aldrich) and 2.65 mg/mL of PECTINASE (17389, Sigma-Aldrich) in 50 mM sodium acetate buffer (pH 5.0) at 50 °C for 72 h under continuous inversion using a mini-labroller (Labnet, USA) placed in an INCU-line IL23 (VWR, Sweden). Following incubation, samples were centrifuged at 10 000 × g for 10 min and the supernatant was removed. The remaining cell wall pellet was isolated as described above^15^ and air-dried overnight. Enzymatically isolated lignins were stored on the bench in closed microtubes for several months prior to characterization.

### Lignin content and composition determination

Lignin composition was determined using pyrolysis-GC/MS to measure S/G and GCHO/GCH_2_OH on 60 µg (±10 µg) of dried 8-week-old stem samples as previously described^26,31^. Lignin content was measured using the CASA method as previously described^56^: 10 mg of dried extractive-free isolated cell walls placed in microtubes with a safety lock were supplemented with 10% cysteine (168149, Sigma-Aldrich) solution in 70% H_2_SO_4_ solution in water (w:w) and vortexed horizontally for 1h. Samples were centrifuged at 10,000g for 10 min and the supernatant was diluted 1:20, 1:100 and 1:200 in ultrapure water, placed in a UV-compatible 96-well plate and absorbance spectra at 283 nm were measured using multiplate reader (Hidex, Sweden). The percentage of CASA lignin (weight percentage in extractive-free cell walls) was calculated using Eq. 1.

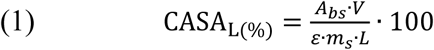

With Abs is the absorbance at 283 nm, V is the volume (L), ε is the absorption coefficient (12 g^−1^·L cm^−1^), m_s_ the mass of cell wall (g) and L is the light path length (0.529 cm).

### UV-Vis-NIR absorbance microspectrometry

UV-Vis-NIR microspectra were obtained using the FLEX PRO UV-Vis-NIR microspectrophotometer (CRAIC Technologies, Optoprim Germany). Samples were point- sampled using the 10x objective (NA 0.20) under AP4 settings (optical resolution ∼25 µm) or AP6 (optical resolution ∼1 µm). Illumination was set to achieve a photon count between 30000 and 50000 in the visual range by setting the standard integration time to 300 (halogen lamp) and 800 (deuterium lamp) µs. Spectra were acquired in transmission mode from 168 to 1200 nm at a spectral resolution of 0.4 nm. Spectral data were exported into ‘csv’ format and further analysed in Matlab (2025b, Mathworks). Upon import into Matlab, deuterium and halogen spectra were combined for reference spectra (ref, no object within the light path), black body spectra (blk, full deflection of light away from the detector) and sample spectra (smpl, transmission through sample and substrate) as absolute photon count (N_p_(λ)) spectra.

Transformation into rel. transmission spectra was done by calculating the photon count difference of the measured spectrum to the black body and reference spectrum according to Eq. 2-5.

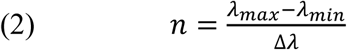

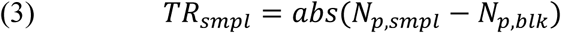

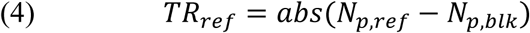

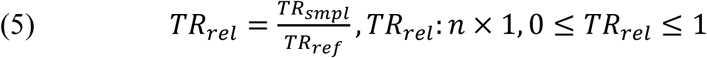

Relative transmission spectra were transformed into absorbance spectra according to Eq. 6.

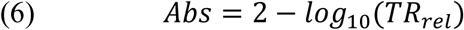

For analyses using principal component analysis (PCA) and pure color determination, spectra were range normalized by setting the minimal absorbance (0.1% quantile) in each spectrum to 0 and the maximal absorbance (99 % quantile) to 1. For range normalization in transmission spectra the opposite operation was conducted: the maximum transmission (minimum absorbance, as described above) is set to 1 and the minimum transmission (maximal absorbance) is set to 0. This procedure normalizes density and illumination effects on color and relative differences within spectra.

### Analysis and color determination

Color was determined using the 10-degree standard observer as described by the International Commission on Illumination (CIE, 1964). Appropriate reference spectra from the CIE repository (https://cie.co.at/datatable/cie-1964-colour-matching-functions-10-degree-observer) was used and converted the wavelength range and resolution to fit our measured spectral range, by restricting the spectral range between 360 and 830 nm and increasing the spectral resolution of the CIE standards from 1 nm to 0.4 nm accordingly, while normalizing the overall area under the curve to 100% for each reference standard.

Color was determined by calculating the XYZ color space using the described spectral reference standards for the X, Y and Z imaginary colors perceived by either the 2- or 10-degree standard observer. The 10-degree standard observer represents a wider field of view as perceived in natural conditions. The 2-degree standard observer is often used in microspectral correction due to the smaller field of view of the observer better representing what would be seen through a microscope. The 10-degree standard is usually applied to standard spectrophotometers as it better describes wide-field vision. In our case we do not intend to mimic the restricted field of view of the microscope, but rather extrapolate the perceived wide field appearance from our spectral information. Therefore, we choose the 10-degree observer and the X, Y and Z color codes were calculated by multiplying the relative transmission spectra with intensities between 0 and 1 by the X, Y and Z reference standard spectra. The color coordinates in the XYZ color space are computed from the area under the curve of the multiplied spectra. The resulting values for XYZ relate to a neutral ideal light source with equal illumination across the entire spectral range. This abstract color space can be transformed into the universal L*a*b* color space using the following equations.

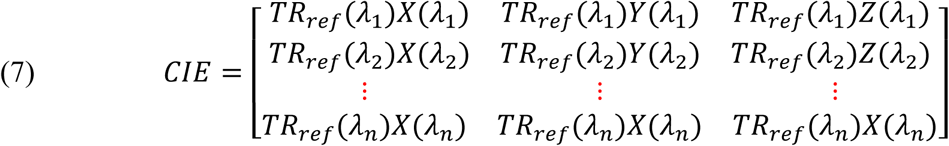

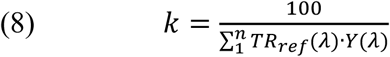

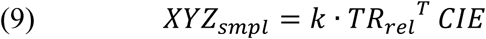

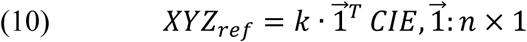

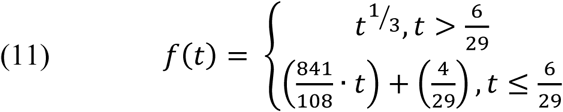

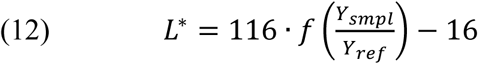

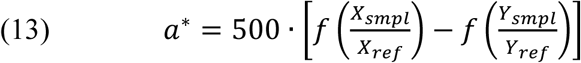

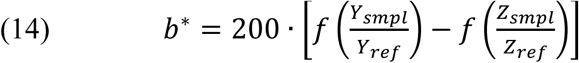

The L*a*b* color space is a numerically independent color space in which L* represents lightness between 0 and 100, with axes a* ranging between ‘green’ (positive) and ‘red’ (negative) and b* ranging between ‘yellow’ (positive) and ‘blue’ (negative). To better relate to traditional color perception, we transformed this colorspace into a polar coordinate system, namely CIE L*C*h°, describing hue as the rotational angle described by Eq. 15 and Chroma (C), similar to saturation, as the radial distance of the hue coordinates Eq. 16 similar to the HSL colorspace.

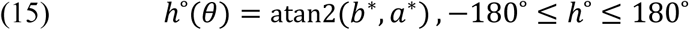

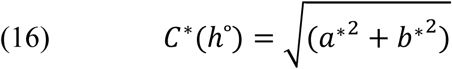

Because standard HSL is tethered to the artificial boundaries of hardware displays (sRGB), it distorts human perceptual realities—such as the fact that the eye perceives yellow as inherently brighter than blue. To bridge intuitive human interpretation with strict colorimetric accuracy, we mapped 3,600 evenly sampled HSL coordinates into the CIE L*C*h° space using an equal-energy Illuminant ’E’ reference and the ’prophoto-rgb’ gamut. The maximum theoretical Chroma (C*) achieved for each hue angle (θ) under this system was defined as the 100% saturation boundary. Consequently, all experimental material colors are reported as a percentage of this absolute perceptual limit C*_sample_(θ) / C*_max_(θ). Unlike the relative saturation index (C*_sample_(θ) / C*_max_(θ), the Lightness L* component was maintained on its absolute colorimetric scale 0 to 100. This preserves the physical reality of human visual perception, where different saturated hues possess inherently different peak luminance levels relative to the equal-energy reference (Illuminant E). Normalizing L* against hue-specific maxima would distort the true perceptual brightness, masking the high luminance of yellows and the low luminance of blues. However due to digital display format standards and reader preference to visualize data on electronic display screens L*a*b* scores were converted to RGB-values using the ‘sRGB’ gamut for graphical displays in this publication, whereas all numerically tabulated data refers to ’prophoto- rgb’ for RGB and HSL conversion. Supplementary Data S2 demonstrates a Matlab workflow, showing the import of CRAIC instruments exported CSV-files of UV-Vis spectra and the subsequent optional normalization and color data extraction.

Statistical assessments were performed either on absorbance based spectral information or L*a*b* / L*C*h° color information. Parameters were compared using non-parametric tests by evaluating groups using KS-testing, quantile absolute distances (QAD), and divergence effect size (D) to evaluate statistical significance. For absorbance spectra groups were evaluated at each wavelength interval independently on individual data per group and wavelength interval, as well as correlatively by calculating the Pearson correlation of group average spectra. Each statistical comparison created vectors for p_ks_(λ), D_ks_(λ) and QAD(λ), enabling the definition of spectral regions related to compared group traits. To determine whether groups were significantly different, we reduced the dimensionality of the test by PostHoc utilization of the False – Discovery- Analysis after Benjamini-Hochberg as well as permutation testing, which computes the critical values D_ks,crit_(λ) for D_ks_(λ) as max(D_ks_(λ)) for 10000 randomized sample groupings. The p_PostHoc, Permutation_ is represented by n_λ_(D_ks_(λ) > D_ks,crit_(λ))/n_λ_.

### Principal Component Analysis (PCA) and hierarchical clustering

Principal component analysis (PCA) was based on the spectral data from synthesized DHPs between 280 and 1000 nm. The resulting Loading vectors were used to define principal component contributions to samples of different origin. This was done to keep dimensionality between different sample groups consistent and reference all data towards this synthetic reference space. Hierarchical clustering analysis (HCA) was performed based on the PCA component scores. Dendrograms were manually reordered to keep the branching order consistently transitioning from light to yellow/orange to red. Both PCA and HCA data did not include calculated L*a*b* data. Thus any colorimetric alignment between these parameters is solely based on spectral properties of the original data after range normalization.

## ACKNOWLEDGEMENTS

This work was supported by the Circular Bio-based Europe Joint Undertaking (CBE-JU) and its members (Project Bio-LUSH, grant no. 101112476 to EP), the Swedish research council Vetenskapsrådet (VR) research grants 2023-03661 (to EP), the Swedish research council for sustainable development FORMAS research grant 2023-00778 (to DM and EP), the Bolin Centre for Climate Research RT2 and RT4 “seed money” (to AG and EP), the Carl Trygger Foundation CTS16:362/CTS17:16/CTS 23:2756 (to EP) and the Knut and Alice Wallenberg Foundation (KAW) through the Wallenberg Wood Science Center (KAW 2021.0313). We particularly thank Stiftelsen Nils och Dorthi Troëdssons forskningsfond for funding to acquire the UV-Vis-NIR microspectrometer (project 1126-2024 to EP). We also thank the Sven och Lilly Lawskis fond för naturvetenskaplig forskning for post-doctoral support (to AG). We thank Ms. Indigo Tazaki and Sarah Boyle for help in preparing *Arabidopsis* stem sections, enzymatically isolating lignins and the CASA stem lignin measurements. We thank Prof. Katharina Pawlowski for critical comments. We thank Dr Nobutaka Mitsuda and Dr. Shingo Sakamoto from the National Institute of Advanced Industrial Science and Technology (AIST, Japan) for supplying seeds of *C4H:FAH1* gain-of-function plants. We also thank the Departments of Ecology, Environment and Plant Sciences (DEEP), the Bolin Centre for Climate Research and the Center for Circular and Sustainable System (SUCCeSS) of Stockholm University (SU) as well as the Institute of Global Innovation Research (GIR) of Tokyo University of Agriculture and Technology (TUAT).

## AUTHOR CONTRIBUTIONS

Conceived and designed the experiments: AG, DM, EP. Produced the material: AG, DM, MN, LM, MHS, SES, SK, EP. Performed the experiments: AG, DM, MN, AC, EP. Analyzed the data: AG, EP. Ensured financial support: EP. Wrote the manuscript: EP. All authors then thoroughly revised and commented on the manuscript.

---

**Figure S1.**
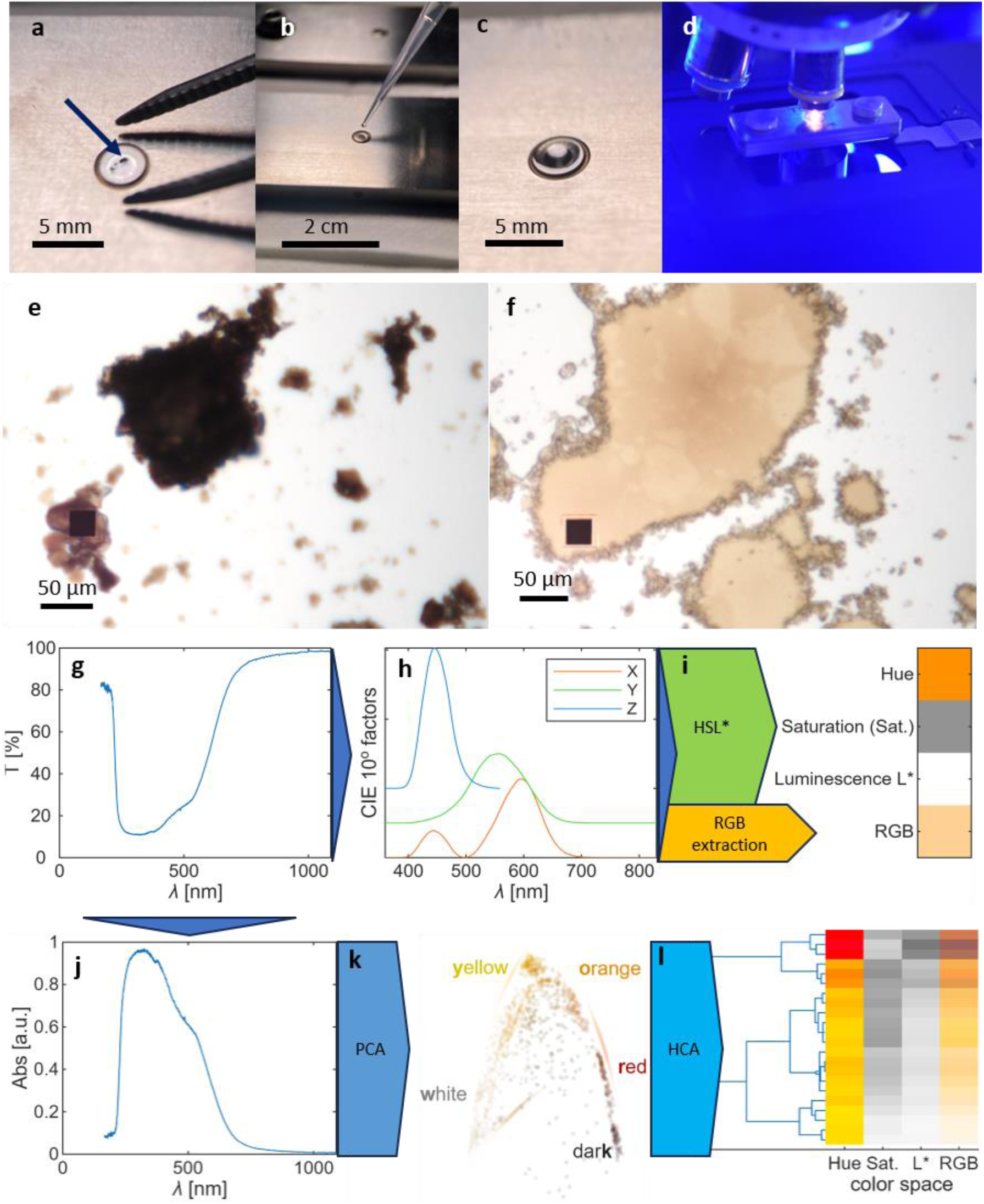
Experimental set-up to reliably measure transmittance/absorbance of insoluble samples using UV-Vis microspectrometry to obtain color information. UV-Vis microspectroscopy can directly be done on tissue samples, isolated fibers/cells, or particle material. Here, we exemplified the use of insoluble synthetic lignins as small particle (blue arrow) directly deposited on the diamond window of the compression cell (a), supplemented with water for hydration (b, c), and compressed to the maximal pressure in the compression cell that is then placed onto the microscope stage for imaging (d). Particles before compression appear dark (e) and after compression become translucent (f). Black squares in (e) and (f) represent the aperture (AP4, 10x, NA = 0.2) used to measure transmittance/absorbance on the mounted samples. Acquired transmission spectra (g) and transformed according to the CIE standard observer spectral weights (h). The resulting XYZ colorspace is then transformed into L*a*b*, a universal color format, using the microscope white reference to obtain correspondence into HSL and RGB color space models (i) using the equal energy radiator reference values. Transmission spectra are also converted into absorbance (j) to undergo data analysis such as principal component analysis (k) and hierarchical clustering (l).

**Figure S2.**
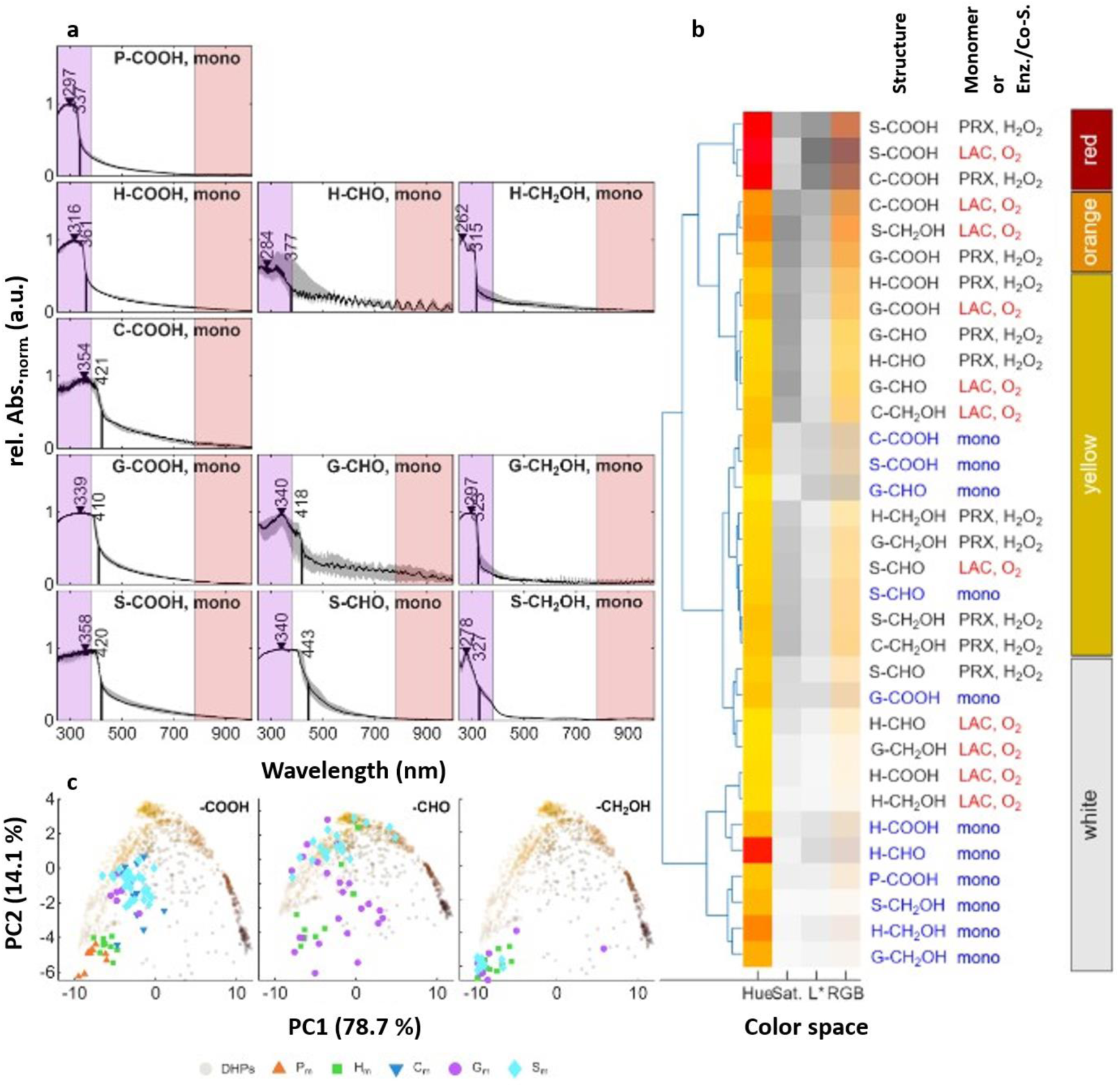
Analyses of monomer absorbances and color information using UV-Vis microspectrometry. (a) UV-Vis-NIR spectra and their IQD (shaded in grey; n = 20-30 particles) of monomeric compounds along the biosynthetic phenylpropanoid pathway varying in ring substitution (either unsubstituted phenyl/P, *para* hydroxylated *p-*hydroxyphenyl/H, *para* hydroxylated *meta* hydroxylated caffeyl/C, *para* hydroxylated *meta* methoxylated guaiacyl/G, *para* hydroxylated *meta* dimethoxylated sinapyl/S) and aliphatic chain functions (either carboxylic acid/COOH, aldehyde/CHO or alcohol CH_2_OH). Spectral peak maxima are indicated by triangle whereas mid-maximal wavelengths towards Vis range are indicated by line. (b) hierarchical clustering analysis of the HSL and RGB color information of unpolymerized monomers compared to their polymerised forms as synthetic lignin DHPs (oxidised using either PEROXIDASE/PRX with H_2_O_2_ or LACCASE/LAC). (c) Principal component analysis (PCA) of unpolymerized monomers compared to their polymerised forms as synthetic lignin DHPs (oxidised using either PEROXIDASE/PRX with H_2_O_2_ or LACCASE/LAC) according principal component (PC) 1 and 2 (see Figure S3).

**Figure S3.**
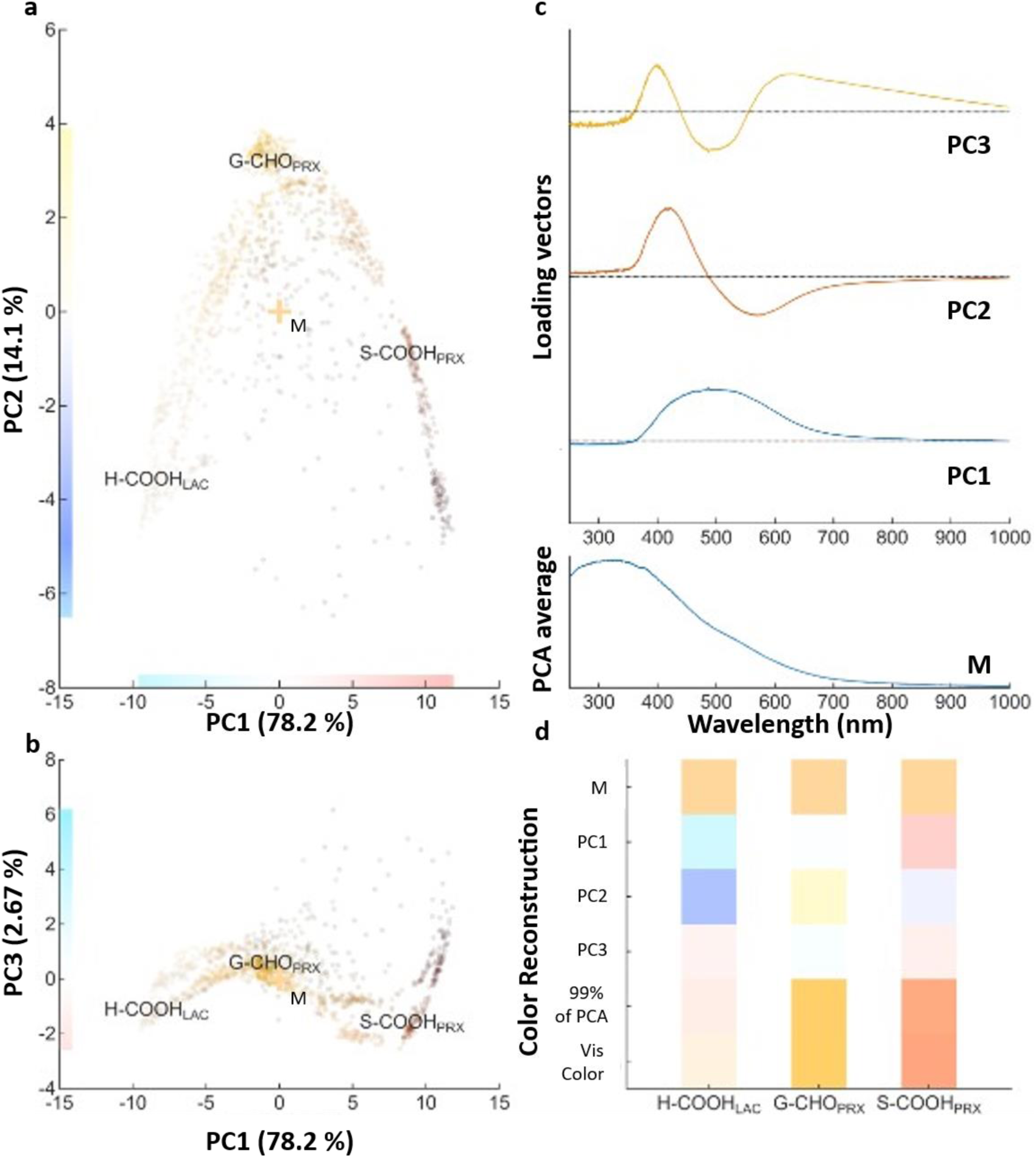
Principal component analysis (PCA) of synthetic lignin homopolymer DHPs. (a, b) the three different principal components (PC) describe specific spectral features of synthetic lignins based on their absorbance. The scattering of the data displays the individual particle spectra which are colored correspondingly to their sRGB color. Three DHPs with extreme properties are indicated onto the scatter plots made of H-COOH (oxidized by LAC, colored white), G-CHO (oxidized by PRX/H_2_O_2_, colored yellow) and S-COOH (oxidized by PRX/H_2_O_2_, colored red) respectively to the PCA mean position (indicated by M). Each axis also includes the respective hue shift from the PCA mean position (M) of each principal component with PC1 explaining 78.1 % of the variance (corresponding to the blue to red spectral shift), with PC2 explaining 14.1% of the variance (corresponding to the blue to yellow spectral shift), and PC3 explaining 2.7% of the variance (corresponding to the rose to turquoise spectral shift). (c) the corresponding loading spectra of each PCs are presented together with the average PCA spectrum corresponding at the mean position (M) along the UV-Vis-NIR range. (d) reconstitution of the chromogenic properties of the three DHPs using the 3 PCs explaining >99 % of variance. The cumulative contribution of the 3 PCs (represented as 99% of PCA) enabled to reconstitute the same hues as the one measured from the spectra (represented as Vis color). The average base hue of all data points (indicated by M_PCA_) was modified accordingly to each PC for each DHP including whitish H-COOH (oxidized by LAC), yellow G-CHO (oxidized by PRX/H_2_O_2_) and red S-COOH (oxidized by PRX/H_2_O_2_).

**Figure S4.**
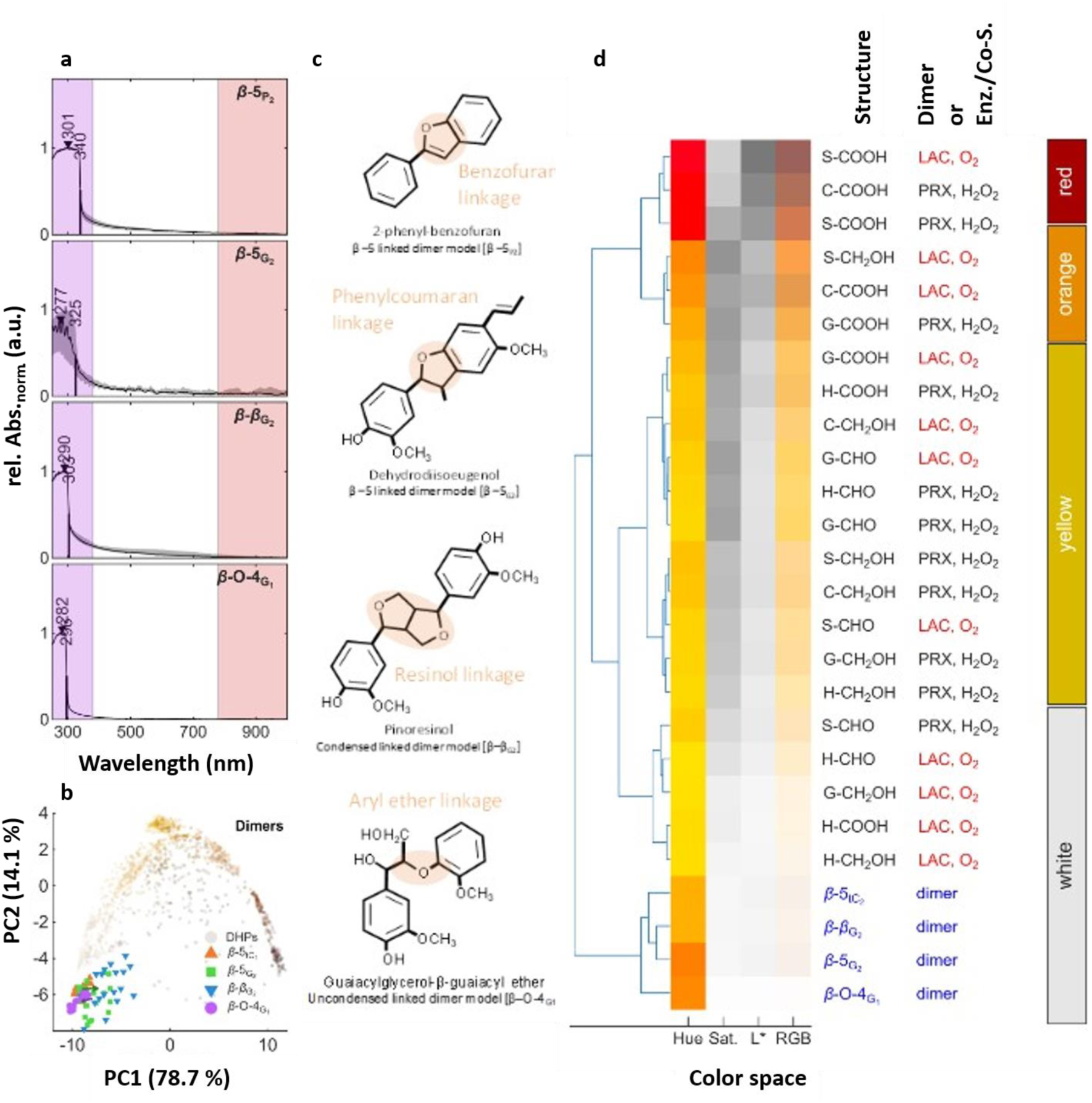
Analyses of dimer absorbances and color information using UV-Vis microspectrometry. (a) UV-Vis-NIR spectra and their IQD (shaded in grey; n = 20-30 particles) of model dimeric compounds varying in interunit linkage type including condensed benzofuran (β-5_P2_), condensed phenylcoumaran (β-5_G2_), condensed pinoresinol (β- β_G2_) and uncondensed aryl ether (β-*O-*4_G1_). Spectral peak maxima are indicated by triangle whereas mid-maximal wavelengths towards Vis range are indicated by line. (b) Principal component analysis (PCA) of model dimers compared to synthetic lignin DHPs (oxidised using either PEROXIDASE/PRX with H_2_O_2_ or LACCASE/LAC) according principal component (PC) 1 and 2 (see Figure S3). (c) Molecular models of the model dimeric compounds used including interunit linkages shaded in orange. (d) hierarchical clustering analysis of the HSL and RGB color information of dimers compared to synthetic lignin DHPs (oxidised using either PEROXIDASE/PRX with H_2_O_2_ or LACCASE/LAC).

**Figure S5.**
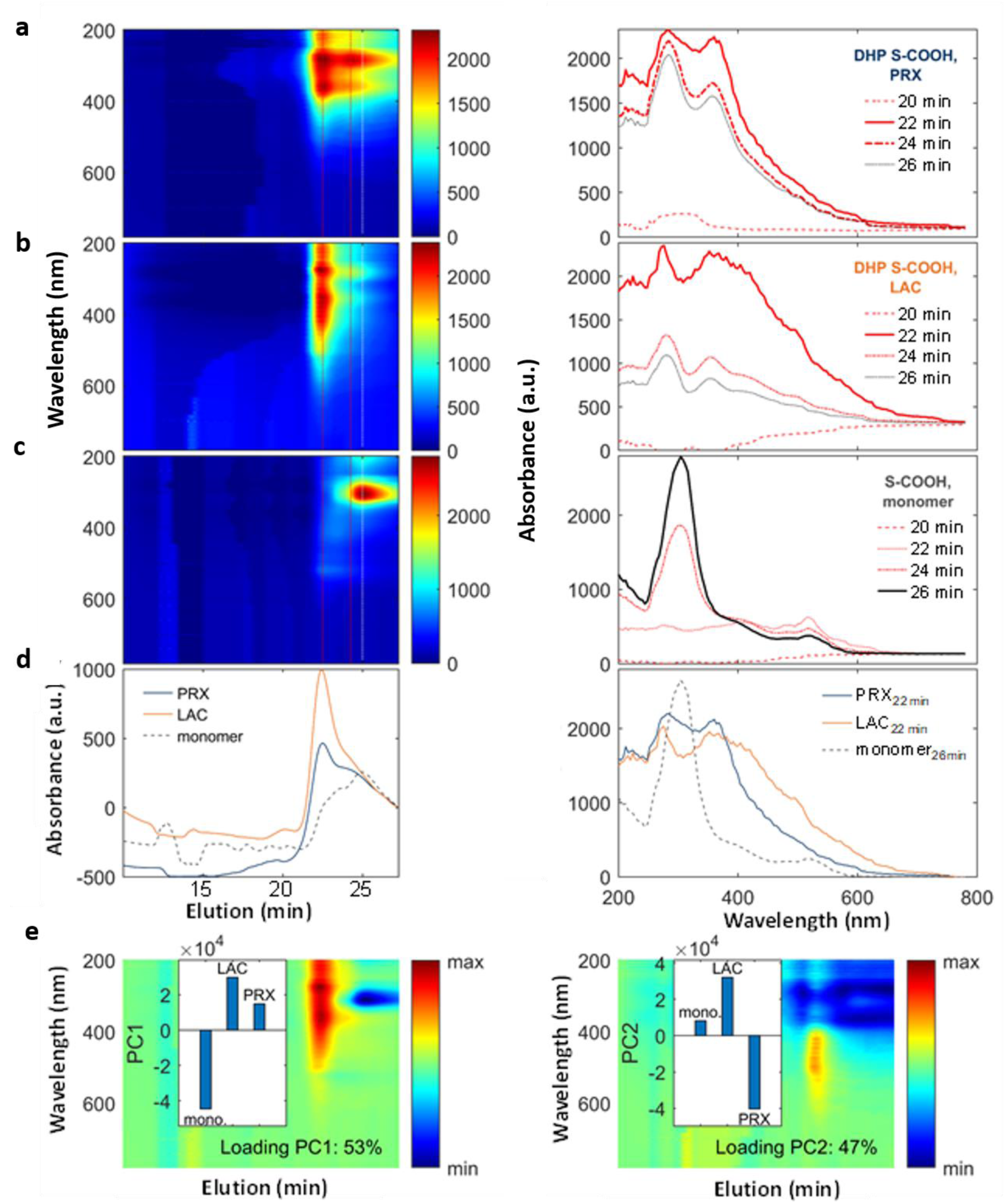
Size exclusion chromatography (SEC) elution profiles of synthetic polymers compared to unpolymerized monomers. (a - d) Multidimensional elution profiles (UV-Vis spectra during elution time) revealing molar weight differences between synthetic lignin DHPs oxidised using either PEROXIDASE/PRX with H_2_O_2_ (a), LACCASE/LAC (b) and unpolymerized monomer S-COOH (c) shown as color intensity coded spectral elution (on the left, a-c) or line plot at 280 nm (d). Corresponding absorbances at different elution times for synthetic lignin DHPs oxidised using either PEROXIDASE/PRX with H_2_O_2_, LACCASE/LAC and unpolymerized monomer S-COOH (on the right, a-d). (e) Principal component analysis (PCA) of the 3 SEC elution profiles including loading of principal component 1 (PC1; e, left), colored coded in relative intensity colors, explaining 53 % of the variance in the elution profile between LAC- and PRX-oxidized synthetic lignin polymers compared to unpolymerized monomers. Principal component 2 (PC2; e, right), colored coded in relative intensity colors, explaining 47% of the variance shows absorbance and molar weight differences between LAC- and PRX-oxidized synthetic lignin polymers.

**Figure S6.**
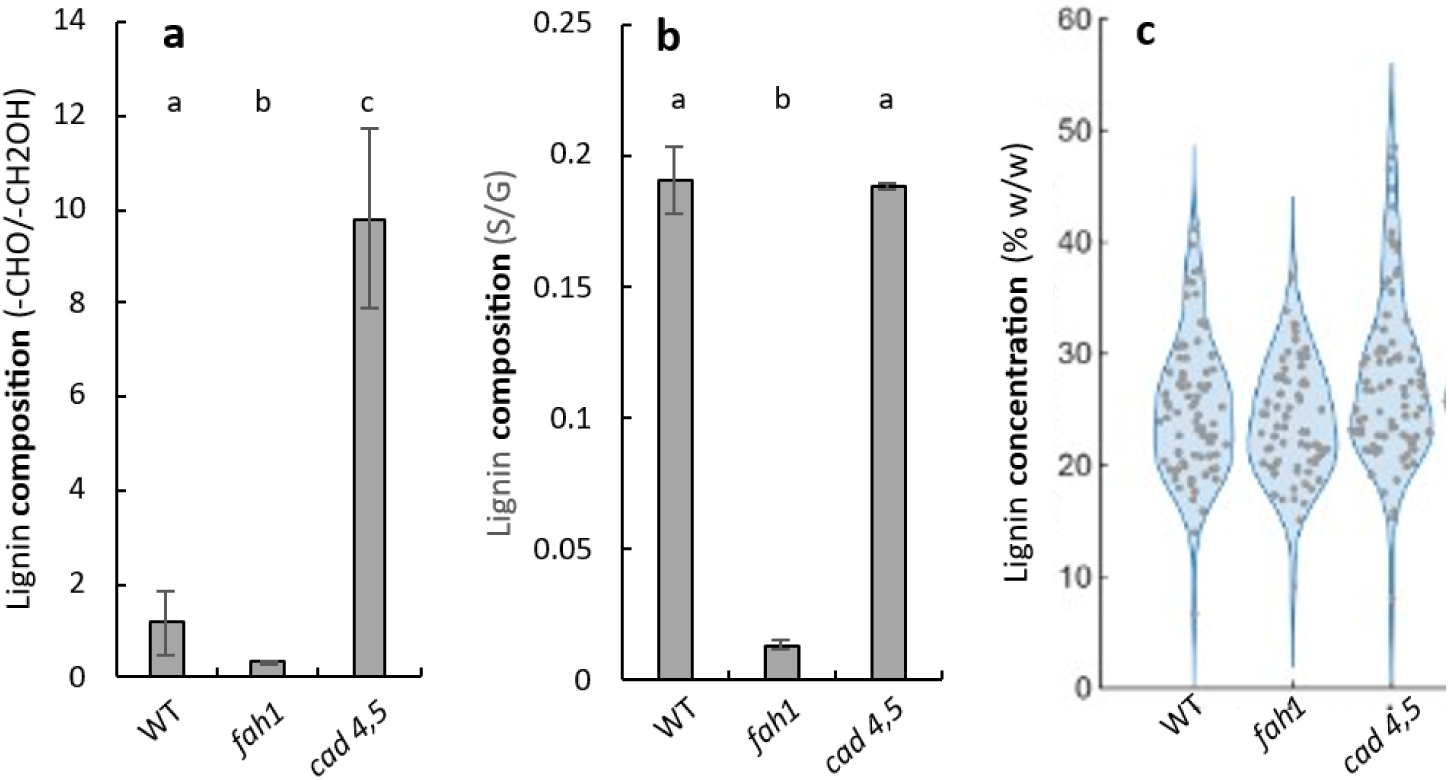
Biochemical analysis of lignin content and unit composition in *Arabidopsis thaliana* wild-type (wt) and loss-of-function mutants in *FERULIC ACID-5-HYDROXYLASE-1* (*fah1*) and in *CINNAMYL ALCOHOL DEHYDROGENASE 4* and *5* (*cad4 cad5*). (a) Bar plots of lignin unit aliphatic aldehyde/CHO to alcohol/CH_2_OH ratio in stems measured using pyrolysis/GC-MS (n = 3-6 individual replicated samples per genotype), bars indicate standard deviation. Different letters indicate significant differences using one-way ANOVA (α = 0.05). (b) Bar plots of lignin unit ring substitution syringyl/S to guaiacyl/G ratio in stems measured using pyrolysis/GC-MS (n = 3-6 individual replicated samples per genotype), bars indicate standard deviation. Different letters indicate significant differences using one-way ANOVA (α = 0.05). (c) Violin plot of cell wall lignin content in stems measured using CASA spectrophotometrical method, expressed as weight percentage of lignin per extractive free cell walls (n = 6 technical measurement of 5 individual replicated samples per genotype).

**Figure S7.**
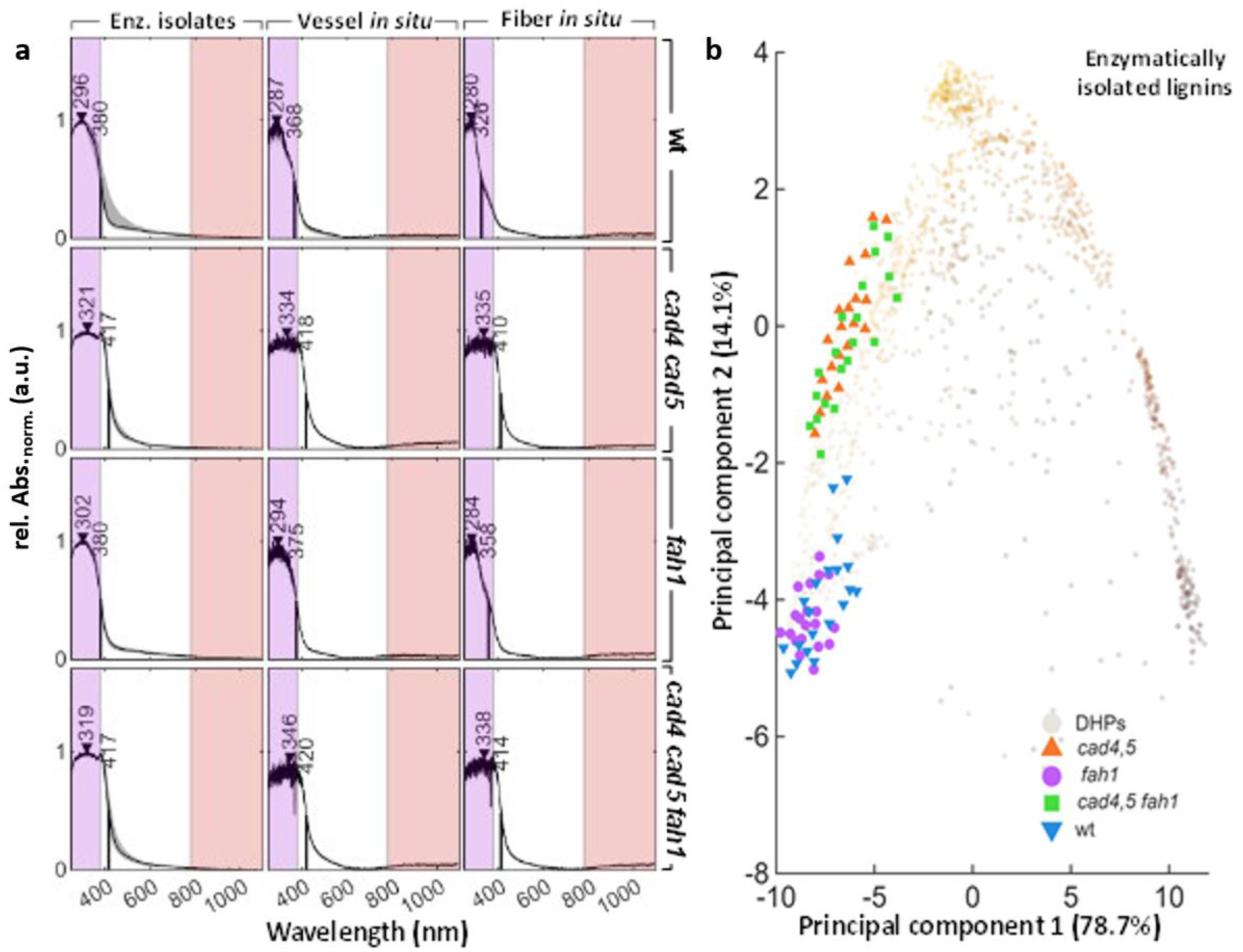
Enzymatic isolated of native lignin. (a) UV-Vis-NIR spectra and their IQD (shaded in grey; n = 20-30 independent samples for particles or cell positions) of enzymatically isolated lignins and lignified cell types (vessels and fibers) measured *in situ* directly in plant biopsies from different *Arabidopsis thaliana* genotypes including wild-type (wt) and loss-of-function mutants in *fah1*, *cad4 cad5*, and *cad4 cad5 fah1*. Spectral peak maxima are indicated by triangle whereas mid-maximal wavelengths towards Vis range are indicated by line. (b) Principal component analysis (PCA) of enzymatically isolated lignins from the different *Arabidopsis thaliana* compared to synthetic lignin DHPs (oxidised using either PEROXIDASE/PRX with H_2_O_2_ or LACCASE/LAC) according to principal component (PC) 1 and 2 (see Figure S3).

**Figure S8.**
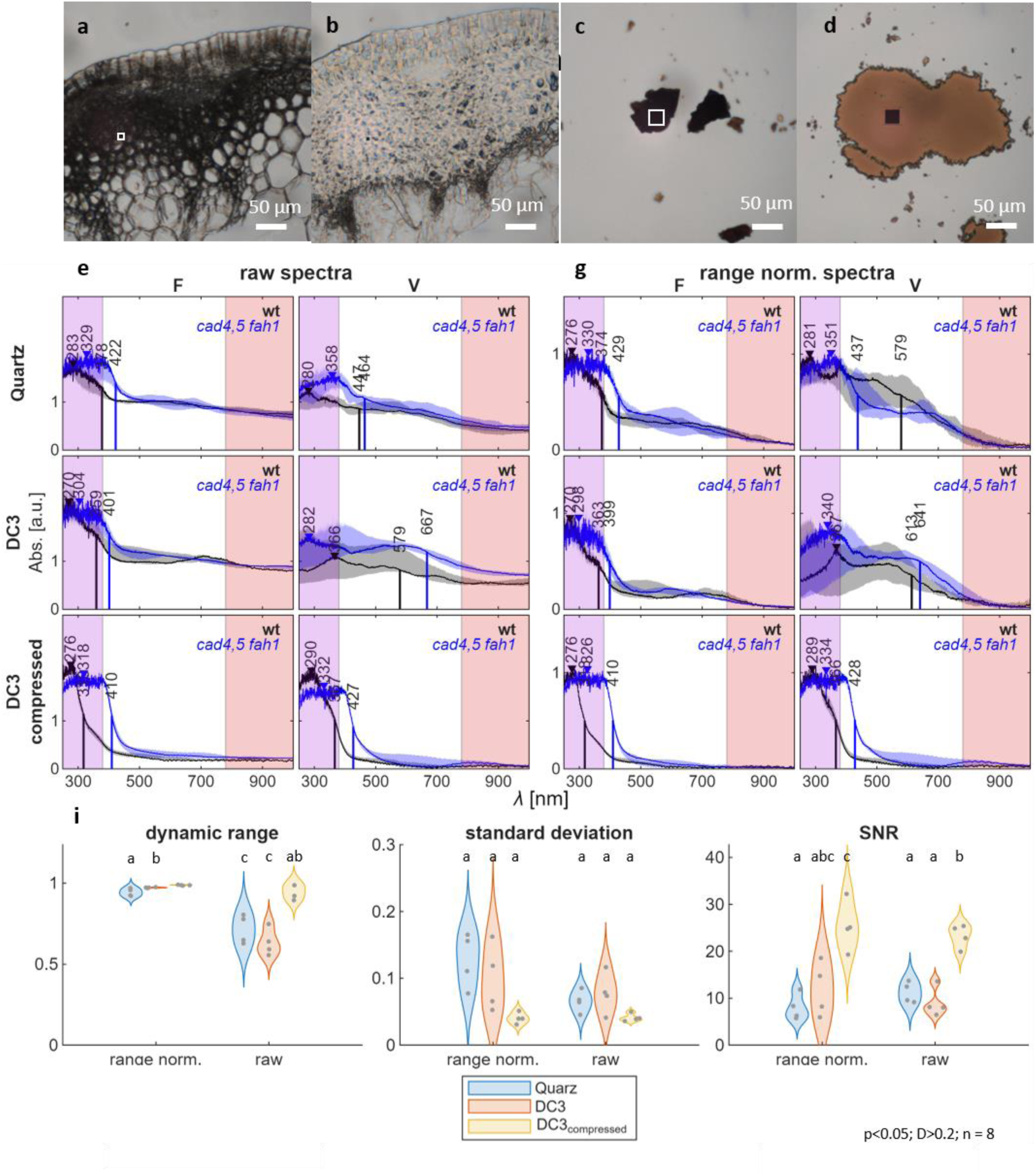
Optimisation of mounting conditions for the *in situ* analysis of plant biopsies using UV-Vis microspectrometry. (a-d) Transmission images of *Arabidopsis* stem cross-section (a-b) and insoluble lignin particles (c-d) before (a, c) and after (b, d) compression in the diamond windowed compression chamber (DC3). Measurement aperture is indicated by white/black boxes. Note that uncompressed samples (a, c) are very dark due to light scattering on irregular phase transitions that are not perpendicular to the light path and/or extend the light path in which light is absorbed, whereas compressed samples (b, d) reducing sample thickness and the number and orientation of phase-transition surfaces, increase the amount of transmitted light and the quality of spectral data. (e, g) Comparison of spectral quality for plant biopsies placed between quartz coverslip and slide, diamond windowed compression chamber without compression (DC3) and with compression (DC3 compressed). Average raw (e) and range-normalised (g) UV-Vis-NIR absorbance spectra and their IQD (shaded in grey; n = 8 cell positions) are provided for fiber (F) and vessel (V) cell types from tissue cross-sections of *Arabidopsis thaliana* wild type (wt) and triple loss-of-function mutant *cad4 cad5 fah1* (labelled *cad4,5 fah1*). (i) Dynamic range, standard deviation and signal-to-noise ratio (SNR) between the mounting conditions are presented. Note the significant effect of the mounting condition on the quality and variation of the spectra measured, with compression allowing for more light to be transmitted/absorbed and less light reflected away from the detector. Statistical differences are indicated with different letter using a one-way ANOVA (α =0.05).

**Figure S9.**
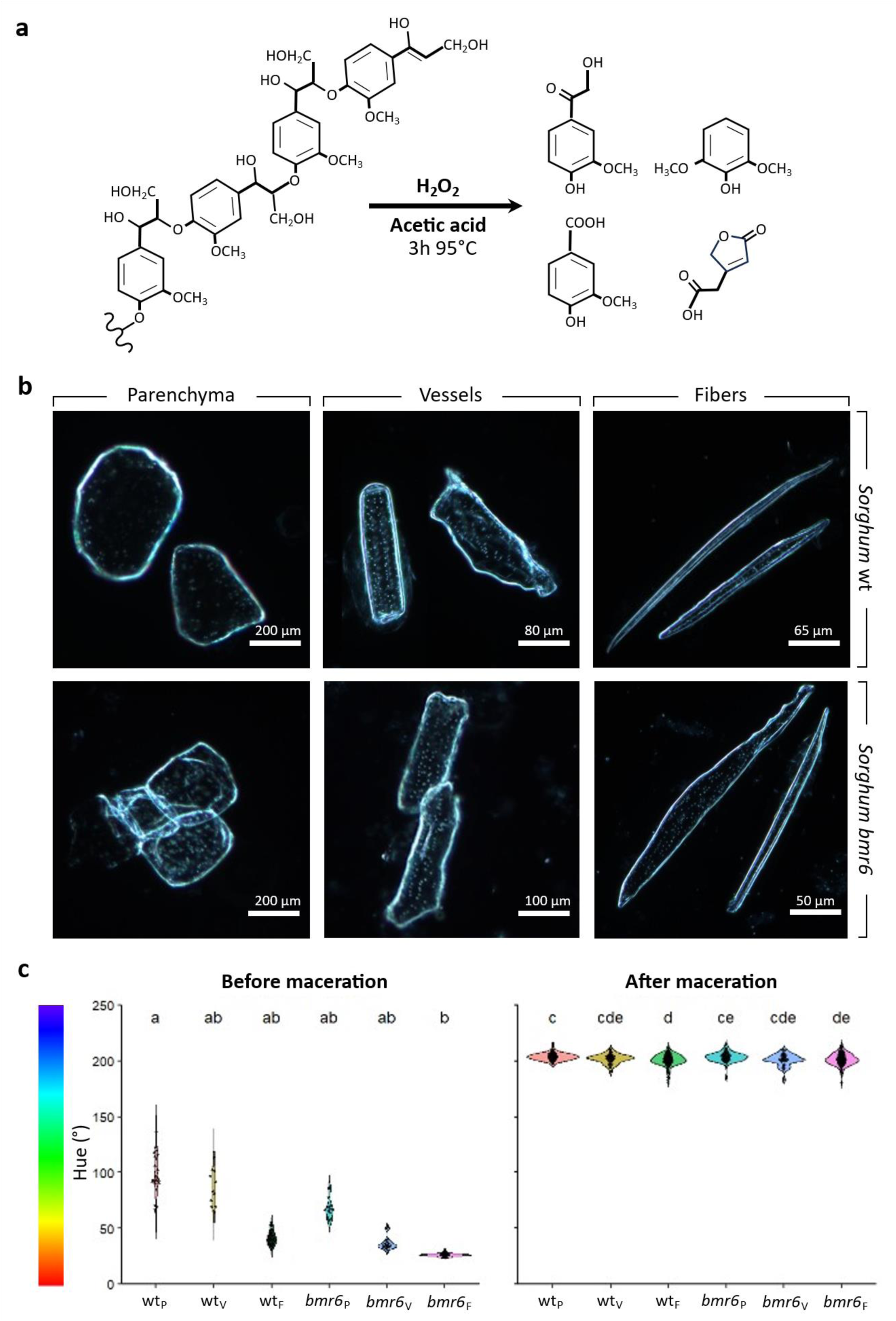
Removal of lignins from wt and *bmr6* sorghum stem samples using oxidation in acid conditions. (a) Schematic representation of oxidative lignin removal during tissue maceration using hydrogen peroxide (H_2_O_2_) in acidic conditions (acetic acid, 95 °C for 3 h)^35–36^. (b) Differential interference contrast (DIC) images of the isolated cell types from wild-type (wt) and *bmr6* sorghum stems after oxidative maceration including parenchyma, vessel and fiber cells. (c) Hue analysis of sorghum parenchyma (P), vessel (V) and fiber (F) cells before and after oxidative maceration, measured on intact stem cross-sections and isolated single cells, respectively. Hue values were quantified using a custom macro for semi-automated detection of regions of interest (ROI) based on intensity thresholding after conversion to an HSB image in Fiji (Supplementary Data S1). Violin plots show the distribution of single-cell measurements. Letters indicate significance determined by a Kruskal-Wallis test followed by Dunn’s multiple comparisons test with Benjamini-Hochberg correction (adjusted *p* < 0.05). A total of 175 cells were analyzed before maceration (n = 11-60 per group) and 472 cells after maceration (n = 32- 136 per group).

**Figure S10.**
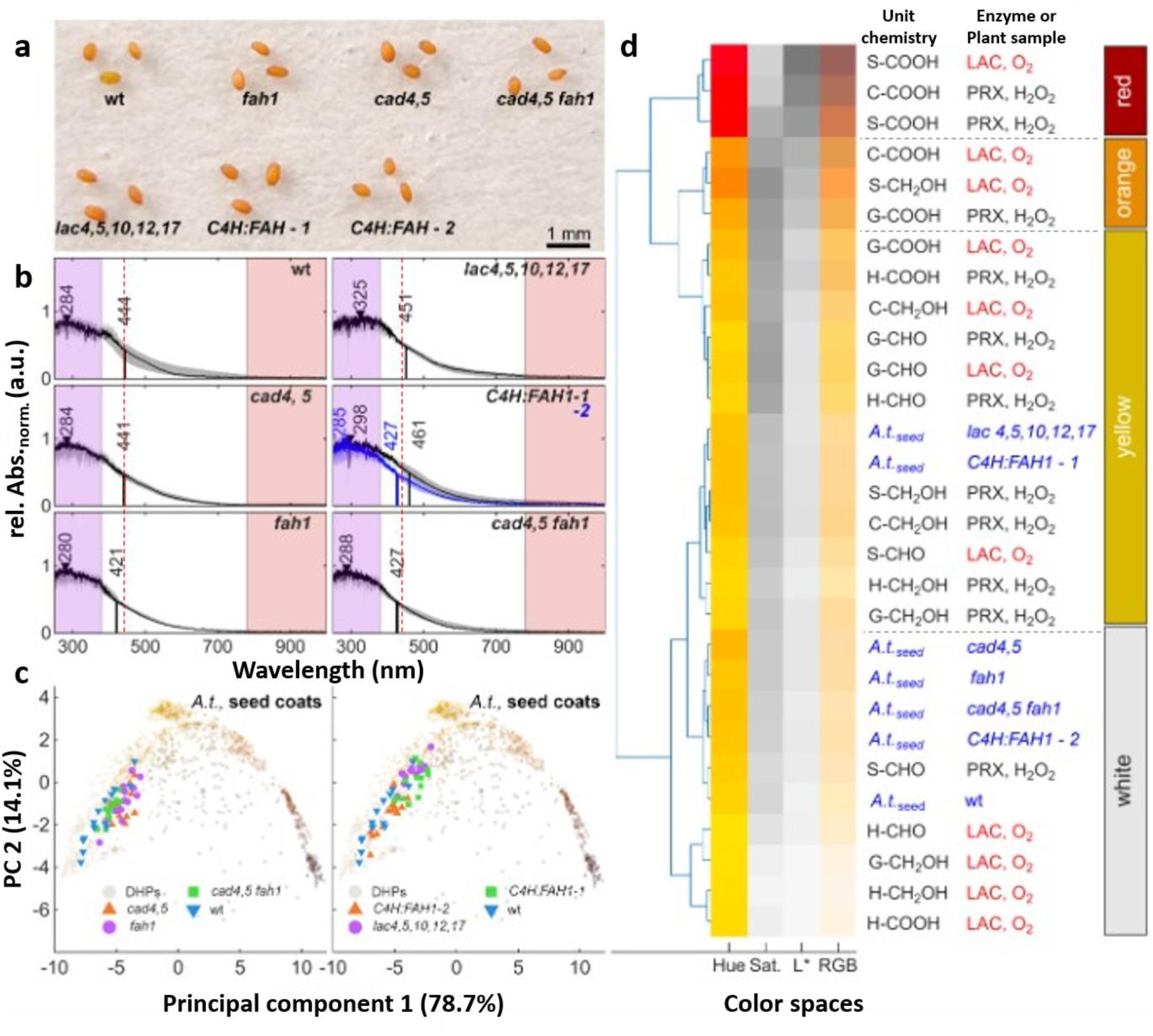
Changing seed coat color using genetic engineering to control the accumulation of specific lignin chromogens. (a) Images of *Arabidopsis thaliana* seeds from different genotypes including wild-type (wt), loss-of-function mutants *fah1 cad4 cad5 cad4 cad5 fah1, lac4,5,10,12,17* as well as gain-of-function mutants overexpressing FAH1 under strong phenylpropanoid-specific promoter (*C4H:FAH1* line 1 and 2). Bar indicates scale. (b) UV-Vis-NIR spectra and their IQD (shaded in grey; n = 20-30 particles) of industrially isolated lignins from different feedstocks (single or mixed species) and different processes including organosolv (OS), sulfate/Kraft, sulfite/lignosulfonate or soda pulping. Maximal absorbances are indicated by triangles and mid-maximal Vis absorbance wavelength with black lines. Note that the large effect of both feedstocks (annuals vs hardwoods vs softwoods) and pulping process on the red-extension of the mid-maximal Vis absorbances. (c) Principal component analysis (PCA) of the absorbance spectra of the different seed coat samples compared to the DHP lignin color palette showing both the color of the different lignins as well as the variability between genotypes. (d) Hierarchical clustering analysis (HCA) of the HSL and RGB color information of the different seed coat lignins compared to synthetic lignin DHPs color palette (oxidised using either PEROXIDASE/PRX with H_2_O_2_ or LACCASE/LAC).

**Figure S11.**
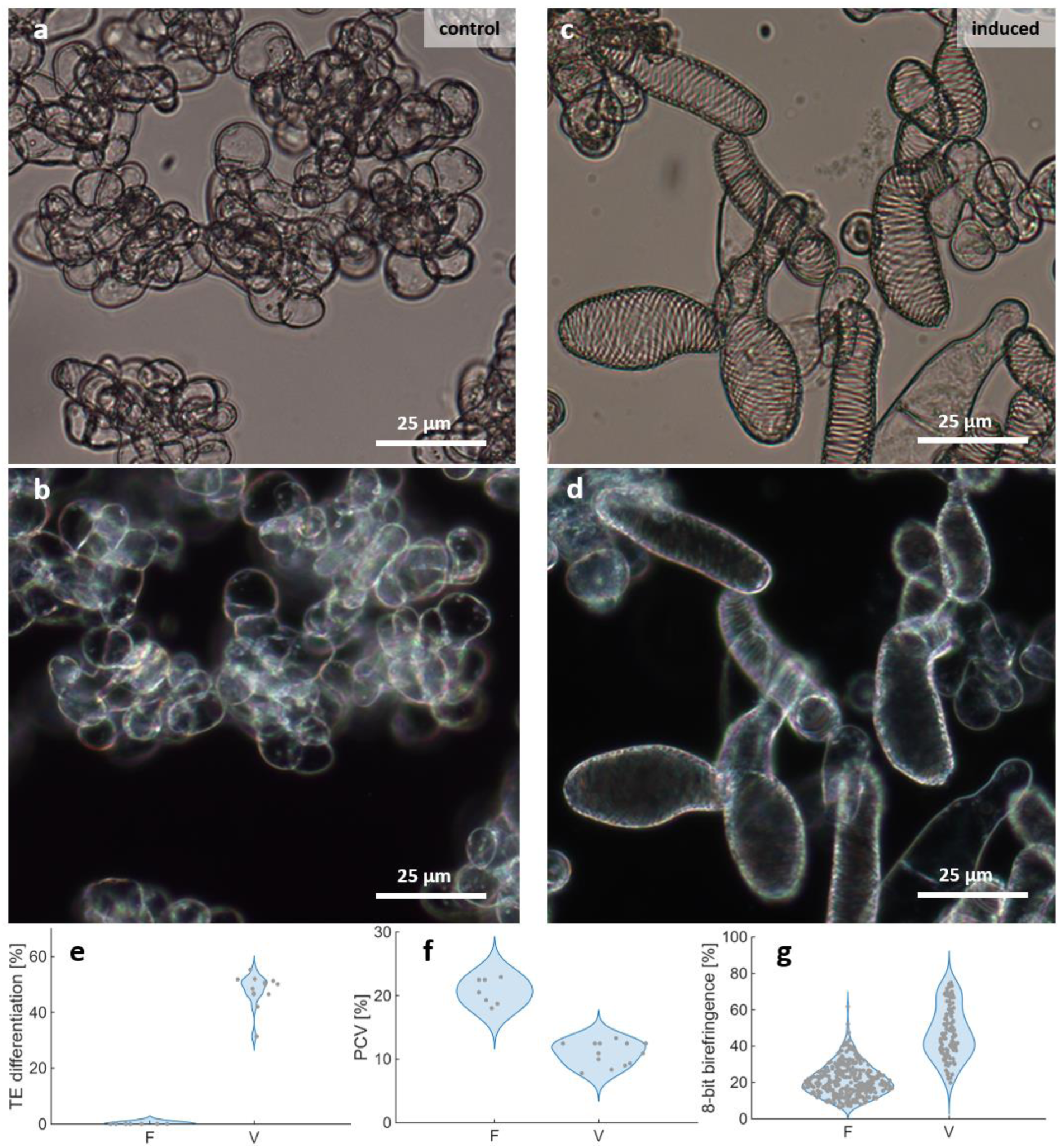
Plant tissue engineering method using plant intercellular tissue cohesion principle through cell walls and not extracellular matrices. (a-d) Brightfield transmission (a,c) and differential interference contrast (DIC; b,d) images of plant inducible pluripotent cell suspensions cultures (iPSCs) triggered hormonally to either differentiate into unlignified parenchymatic cells (a-b) or into lignified wood vessels (c-d). (e) Violin plot of the differentiation efficiency of iPSCs after 7-days of culture after hormonal treatment to induce parenchyma (P) or vessel (V) cells (n = 10-14 independent samples). Note that induced parenchyma cells are completely devoid of vessel cells. (f) Violin plot of the division capacity of iPSCs after 7-days of culture after hormonal treatment to induce parenchyma (P) or vessel (V) cells (n = 10-14 independent samples) expressed as percentage packed cell volume (PCV) after 200g centrifugation per volume of cell suspension culture. Note that induced parenchyma cells are actively dividing in contrast to vessel cells. (g) Violin plot of relative cell wall material deposition per cell by measuring the birefringence of cells in differential interference contrast (DIC) after 7-days of culture after hormonal treatment to induce parenchyma (P) or vessel (V) cells (n = 120-160 independent cells). Note that vessel cells are more birefringent due to their patterned secondary cell walls than parenchyma cells.

**Figure S12.**
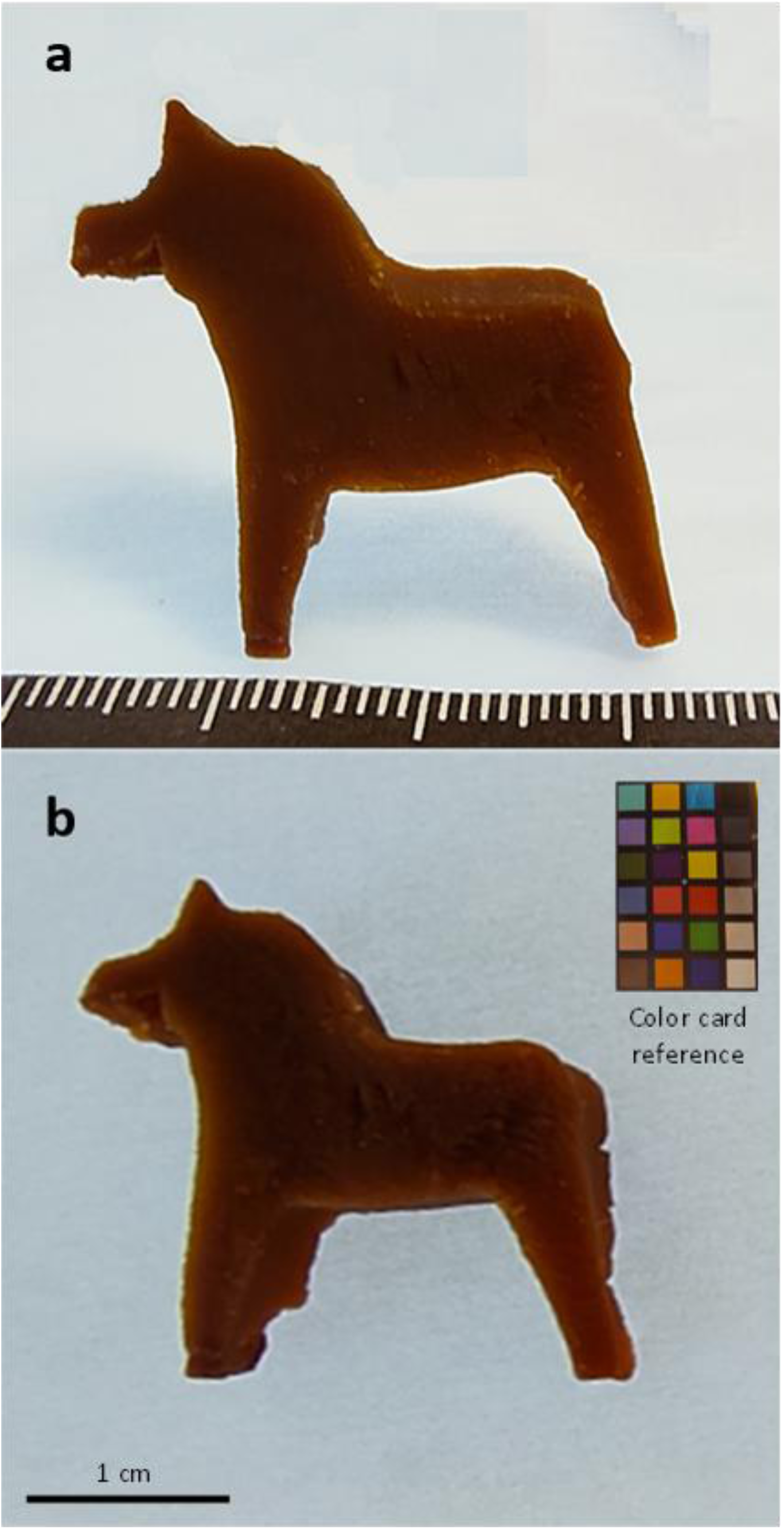
Example of cm wide casting of wood engineered plant tissue into traditional Swedish *dalahäst* (Dalecarlian horse) assembled from vessel-triggered cell types made using inducible pluripotent cell suspension cultures (iPSCs) bound together with externally supplied S-COOH and H_2_O_2_.

**Table S1.** Average range-normalised absorbance spectra of monomeric, dimeric and oligomeric Lignin model compounds.

**Table S2.** Biochemical analysis of the technical lignins used including a The ratio was calculated from the published values1; the ratio itself was not explicitly reported. Condensed phenolic OH includes the integrated condensed phenolic region as defined in the corresponding 31P NMR analysis. The ratio was calculated as condensed phenolic OH (mmol/g) / total phenolic OH (mmol/g). Total phenolic OH includes condensed OH, G-type OH, and H-type OH.

**Supplementary Data S1** Macro for semi-automated cell detection and hue analysis.ijm

**Supplementary Data S2** Functions to import CSV-file exports from LambdaFire (CRAIC technolpgies), Extraction of quantitative and relative cholorimetric values according to CIE defined color spaces (L*a*b*, XYZ, RGB, HSV), as well as a Matlab script and exemplary data to showcase a sample processing pipeline of loading, transforming (Transmission to Absorbance), normalizing and subsequently extracting cholorimatric parameters. Script is tested in Matlab v2025b.

