## Supplementary Data S1 for "Identifying and engineering the molecular origins of lignin color for predictive staining of plant tissues": Supplementary Data S1. Macro for semi-automated cell detection and hue analysis.ijm

```
// ==========================================================
// Supplementary Data 1
// ImageJ/Fiji macro for semi-automated cell detection and hue analysis
// ==========================================================

// CLEAN 
run("Close All");   
roiManager("Reset");
run("Select None");

// OPEN IMAGE
open();
origID = getImageID();

// ROI DETECTION 
run("Duplicate...", "title=mask");
selectWindow("mask");
run("8-bit");
setThreshold(20, 255); //Change if necessary 
run("Convert to Mask");
run("Analyze Particles...", "size=1000-Infinity add");

// CHECKPOINT
roiManager("Show All");
waitForUser("Check ROIs",
"Edit ROIs if needed.\nClick OK to continue.");

// CLOSE MASK
selectWindow("mask");
close();

// ANALYSIS
selectImage(origID);
run("Stack to RGB");
run("HSB Stack");
setSlice(1);

roiManager("Show All");
roiManager("Select All");

run("Select None");   
roiManager("Measure");
```
