## Supplementary Data S2 for "Identifying and engineering the molecular origins of lignin color for predictive staining of plant tissues": importCraicSpecfile.m

```
function UVvisTrans = importCraicSpecfile(filename, dataLines)
%IMPORTFILE Import data from a text file
%  DC3H2OGCHOHPERT018NORM = IMPORTFILE(FILENAME) reads data from text
%  file FILENAME for the default selection.  Returns the data as a table.
%
%  DC3H2OGCHOHPERT018NORM = IMPORTFILE(FILE, DATALINES) reads data for
%  the specified row interval(s) of text file FILENAME. Specify
%  DATALINES as a positive scalar integer or a N-by-2 array of positive
%  scalar integers for dis-contiguous row intervals.
%
%  Example:
% dc3h2oGchohpert018norm = importfile("E:\CRAIC\G-choVchoh\dc3 h2o Gchoh per t0@18_norm.CSV", [2, Inf]);
%
%  See also READTABLE.
%
% Auto-generated by MATLAB on 05-Oct-2025 09:59:45

%% Input handling
% filename = "E:\CRAIC\G-choVchoh\dc3 h2o Gchoh per t0@18_norm.CSV";
% % If dataLines is not specified, define defaults
% dataLines = [2, Inf];

if nargin < 2
    dataLines = [2, Inf];
end

%% Set up the Import Options and import the data
opts = delimitedTextImportOptions("NumVariables", 61);

% Specify range and delimiter
opts.DataLines = dataLines;
opts.Delimiter = "\t";

% Specify column names and types
opts.VariableNames = ["MeasName", "CaseName", "UserName", "Objective", "Aperture", "ImageLoc", "MeasType", "nDateTime", "nDateTimeDarkRef", "CropLow", "CropHigh", "WaveMerge", "MeasMerge", "IntUV", "IntVis", "IntNIR1", "IntNIR2", "AvgUV", "AvgVis", "AvgNIR1", "AvgNIR2", "ResFactVis", "ResFactNIR1", "ResFactNIR2", "MergeUVLow", "MergeUVHigh", "MergeNIR1Low", "MergeNIR1High", "MergeNIR2Low", "MergeNIR2High", "RawMeasUV", "RawMeasVis", "RawMeasNIR1", "RawMeasNIR2", "RawDarkUV", "WaveNIR2", "WaveNIR1", "WaveVis", "MeasUV", "MeasVis", "MeasNIR1", "MeasNIR2", "RawDarkVis", "RawDarkNIR1", "RawDarkNIR2", "RawRefUV", "RawRefVis", "RawRefNIR1", "RawRefNIR2", "Math", "ResponseCorrection", "UVoffset", "NIR1offset", "NIR2offset", "RawRefmerge", "RawDarkmerge", "X", "Y", "Z", "Comment", "Source"];
opts.VariableTypes = ["string", "string", "string", "double", "double", "string", "string", "datetime", "datetime", "double", "double", "double", "double", "double", "double", "double", "double", "double", "double", "double", "double", "double", "double", "double", "double", "double", "double", "double", "double", "double", "double", "double", "double", "double", "double", "double", "double", "double", "double", "double", "double", "double", "double", "double", "double", "double", "double", "double", "double", "double", "double", "double", "double", "double", "double", "double", "double", "double", "double", "double", "double"];

% Specify file level properties
opts.ExtraColumnsRule = "ignore";
opts.EmptyLineRule = "read";

% Specify variable properties
opts = setvaropts(opts, ["MeasName", "CaseName", "UserName", "ImageLoc", "MeasType"], "WhitespaceRule", "preserve");
opts = setvaropts(opts, ["MeasName", "CaseName", "UserName", "ImageLoc", "MeasType"], "EmptyFieldRule", "auto");
opts = setvaropts(opts, "nDateTime", "InputFormat", "dd/MM/yyyy hh:mm:ss aa");
opts = setvaropts(opts, "nDateTimeDarkRef", "InputFormat", "dd/MM/yyyy hh:mm:ss aa");
opts = setvaropts(opts, ["Objective", "Aperture"], "TrimNonNumeric", true);
opts = setvaropts(opts, ["Objective", "Aperture"], "ThousandsSeparator", ",");

% Import the data
data = readtable(filename, opts);
UVvisTrans.SpecUVvisRaw.RawMeasUV = data.RawMeasUV;
UVvisTrans.SpecUVvisRaw.RawMeasVis = data.RawMeasVis;
if mean(isnan(data.RawMeasVis))>0.5
    UVvisTrans.SpecUVvisRaw.RawMeasVis = data.MeasMerge;
end
UVvisTrans.SpecBlkRaw.RawDarkUV = data.RawDarkUV;
UVvisTrans.SpecBlkRaw.RawDarkVis = data.RawDarkVis;
if mean(isnan(data.RawDarkVis))>0.5
    UVvisTrans.SpecBlkRaw.RawDarkVis = data.RawDarkmerge;
end
UVvisTrans.SpecRefRaw.RawRefUV = data.RawRefUV;
UVvisTrans.SpecRefRaw.RawRefVis = data.RawRefVis;
if mean(isnan(data.RawRefVis))>0.5
    UVvisTrans.SpecRefRaw.RawRefVis = data.RawRefmerge;
end
UVvisTrans.SpecWL.WaveVis = data.WaveVis;
if mean(isnan( data.WaveVis))>0.5
    UVvisTrans.SpecWL.WaveVis = data.WaveMerge;
end
end
```
