## Supplementary Data S2 for "Identifying and engineering the molecular origins of lignin color for predictive staining of plant tissues": importUVvis.m

```
function [absor,trans,wavelength,lamp,UVvisTrans] = importUVvis(file)
    %% import CRAIC spectral file
    % file = 'dc3 h2o G-CHO perox l@1.csv';
    UVvisTrans = importCraicSpecfile(file);
    
    countSmpl = [cat(1,UVvisTrans.SpecUVvisRaw.RawMeasUV),...
        cat(1,UVvisTrans.SpecUVvisRaw.RawMeasVis)];
    
    countBlk = [cat(1,UVvisTrans.SpecBlkRaw.RawDarkUV),...
        cat(1,UVvisTrans.SpecBlkRaw.RawDarkVis)];
    
    countRef = [cat(1,UVvisTrans.SpecRefRaw.RawRefUV),...
        cat(1,UVvisTrans.SpecRefRaw.RawRefVis)];
    
    wavelength = cat(1,UVvisTrans.SpecWL.WaveVis);

    %% [Optional] Take out artifact of measurement created by lamp emission spike. 
    % This may not be necessary for your purposes.
    uvTakeOut = wavelength<650|wavelength>660;
    
    countSmpl(:,1) = interp1(wavelength(uvTakeOut),countSmpl(uvTakeOut,1),wavelength);
    countBlk(:,1) = interp1(wavelength(uvTakeOut),countBlk(uvTakeOut,1),wavelength);
    countRef(:,1) = interp1(wavelength(uvTakeOut),countRef(uvTakeOut,1),wavelength);
    
    %% [Optional] Check if spectrum is not just noise. Will set black body spectrum to 
    % 0 if both sample and blackbody spectrum area too close together by 
    % their means (5%).
    SoB = [mean(countSmpl);
        mean(countBlk)];
    SoB = SoB(1,2)./SoB(2,2);
    if SoB<1.05
        countBlk = countBlk.*0;
    end
    %% create transmission spectrum
    countRefs = countRef;                   % light source [UVlamp, Vislamp]
    countRefs(:,:,2) = countBlk;            % black body in 3rd dimension
    %%% absolute values
    %%%% sum under curve
    countNet = sqrt(sum(countSmpl-countRefs,2,'omitnan').^2);
    %%%% spectra
    countNetUVvis = sqrt((countSmpl-countRefs).^2);
    
    %%%% absolute transmission spectrum [UV lamp contribution, VIS lamp
    %%%% contribution]
    countNet = permute(countNet,[1,3,2]);
    %%%% relative to 100% transmission
    transRel = countNet(:,2)./sum(countNet,2,'omitnan');
    transRelUVvis = countNetUVvis(:,:,2)./sum(countNetUVvis,3,'omitnan');
    %% create 
    trans = [transRel,transRelUVvis];
    absor = -log10(trans);
    lamp = [sum(countRefs,2),countRefs];
    lamp = abs(diff(lamp,[],3));
end
```
