## Supplementary Data S2 for "Identifying and engineering the molecular origins of lignin color for predictive staining of plant tissues": spec2cie.m

```
function [Lab,hueD,qSat,SphLab,rgb,hsv_,tbl] = spec2cie(transRel,wavelength,varargin)
    %%%% Info %%%%%%%%%%%%%%%%%%%%%%%%%%%%%%%%%%%%%%%%%%%%%%%%%%%%%%%%%%%%%%%%%
    % This function translates a transmission spectrum into the CIE XYZ color
    % space. From there it computes the L*a*b* color coordinates. These can be
    % converted to hsv or rgb using standard matlab functions.
    %%%% input %%%%%%%%%%%%%%%%%%%%%%%%%%%%%%%%%%%%%%%%%%%%%%%%%%%%%%%%%%%%%%%%
    % transRel = transmission spectrum wavelength x 1 (vector)
    % transRel should be between 0 and 1.
    % wavelength = number of wavelength x 1 (vector)
    % varargin:
    % lamp is an optional variable created from the first input in varargin and
    % represents the white point of the refered spectrum, aka. transmission at
    % 100 %
    %%%% output
    %%%% %%%%%%%%%%%%%%%%%%%%%%%%%%%%%%%%%%%%%%%%%%%%%%%%%%%%%%%%%%%%%%%%%%%%%%
    % Lab = [L*, a*, b*] ... L*a*b* color space defined by CIE
    % hueD ... hue in degrees from 0 (magenta) over red, yellow, green to
    %           180/-180 (cyan), blue, purple to 360/0 (magenta).
    % qSat = [Sample, Gamute Edge] ... absolute Saturation / Chroma in L*
    %           a* b* space
    % SphLab = [hue(rad) or az, elevation (rad), distance from black,
    %           distance from black of gamut edge]
    % rgb = [red, green, blue] ... rgb code for prophoto-rgb gamut (larger
    %           than sRGB), for 'sRGB' values use rgb = lab2rgb(Lab,"WhitePoint","e");
    % hsv_ = [hue, saturation, value] ... for rgb in prophoto-rgb gamut
    % tbl ... all essetial variables in 1 table (can be exported to excel
    
    %%% check for optional 100% spectrum %%%%%%%%%%%%%%%%%%%%%%%%%%%%%%%%%%%%%%
    try
        lamp = varargin{1};
        %normSun = false;
    catch
        lamp = ones(size(wavelength));
        %normSun = false;
    end
    useG = true;
    
    %% % load standard observer CIE spectra %%%%%%%%%%%%%%%%%%%%%%%%%%%%%%%%%%%%
    obsDir = 'Lib\HumanStandardObserver\';
    obsFile = 'stdOb10.mat';
    load(fullfile(obsDir,obsFile));
    
    %% % create X Y Z laodings to fit wavelength
    wl10 = stdOb(:,1);      % wavelength of standard observer
    X10 = stdOb(:,2);       % loading for X
    Y10 = stdOb(:,3);       % loading for Y
    Z10 = stdOb(:,4);       % loading for Z
    %%%%% convert to wavelength
    dWl = mean(diff(wavelength))./mean(diff(wl10));     % X10, Y10 and Z10 are 
    % scaled to sum() = 100. dWl rescales the weights of each wavelength in
    % x10, y10 and z10 to also satisfy: sum() = 100.
    x10 = interp1(wl10,X10,wavelength).*dWl;
    y10 = interp1(wl10,Y10,wavelength).*dWl;
    z10 = interp1(wl10,Z10,wavelength).*dWl;
    % %%%%% check vector sums
    % wlSee = sum([x10,y10,z10],2,'omitnan');
    % wlSee = cumsum(wlSee);
    % wlSee = wlSee./max(wlSee);
    % idxSee = [find(wlSee==0,1,'last'),find(wlSee==1,1,'first')];
    % wlSee(1:idxSee(1)) = 0;
    % wlSee(idxSee(2):end) = 0;
    % wlSee(idxSee(1):idxSee(2)) = 1;
    
    %%% get XYZ colorspace %%%%%%%%%%%%%%%%%%%%%%%%%%%%%%%%%%%%%%%%%%%%%%%%%%%%
    % vector conversion to reference white spectrum:
    S = lamp.*[x10,y10,z10];    
    S(isnan(S)) = 0;
    % transRel clean up
    v = reshape(transRel,1,[]);
    v(isnan(v)) = 0;
    % correction factor for reference white intensity
    k = 100./sum(lamp.*y10,'omitnan');
    % XYZ color space for CIE observer 10 degree
    XYZ = k*(v*S);                          % transRel
    XYZSat = k*(ones(size(v))*S);           % white point
    
    %%% convert to L*a*b* %%%%%%%%%%%%%%%%%%%%%%%%%%%%%%%%%%%%%%%%%%%%%%%%%%%%%
    % create relative x y z to white point reference
    yYn = XYZ(:,2)./XYZSat(:,2);
    xXn = XYZ(:,1)./XYZSat(:,1);
    zZn = XYZ(:,3)./XYZSat(:,3);
    
    % compute L*, a*, b*
    L = 116.*labFt(yYn)-16;
    a = 500.*(labFt(xXn) - labFt(yYn));
    b = 200.*(labFt(yYn) - labFt(zZn));
    Lab = [L,a,b];
    
    %% % Done. Everything else can be used to analyze color further %%%%%%%%%%%
    %%% create the hsv reference color space from the prophoto-rgb gamut and
    %%% convert it to L*a*b*.
    rgbRef = hsv(3600);
    LabRef = rgb2lab(rgbRef,"ColorSpace",'prophoto-rgb',"WhitePoint","e");
    % alternate gamuts: "adobe-rgb-1998" or 'sRGB' (default)
    
    % convert Lab values to polar coordinates
    [az,el,r] = cart2sph(LabRef(:,2),LabRef(:,3),LabRef(:,1));
    % ax = Hue
    % el = elevation angle from Black to white
    % r = sqrt(Chroma^2 + L*^2)
    AzElRref = [az,el,r];
    
    [az,el,r] = cart2sph(a,b,L);
    az(isnan(az)) = 0;
    el(isnan(el)) = 0;
    r(isnan(r)) = 0;
    
    %%% find best fitting saturated reference color
    if std([az,el,r])>0
        % calculate the difference between sample and each reference
        azRef = sqrt((AzElRref(:,1)-az).^2);
        % find the closest match
        azRef = find(azRef == min(azRef));
        % select reference color
        AzElRref = AzElRref(azRef,:);
        LabRef = LabRef(azRef,:);
        rgbRef = rgbRef(azRef,:);
        hsvRef = rgb2hsv(rgbRef);
    else
        AzElRref = [0,0,100];
        LabRef = [100,0,0];
        rgbRef = [1,1,1];
    end
    %%% summerize
    % create spherical colorspace values
    rmax = AzElRref(:,3);
    SphLab = [az,el,r,rmax];
    
    % create rgb and hsv values
    rgb = lab2rgb(Lab,"ColorSpace",'prophoto-rgb',"WhitePoint","e");
    hsv_ = rgb2hsv(rgb);
    hueD = rad2deg(SphLab(:,1));                % Hue in degrees
    qSat = [cos(el).*r,cos(AzElRref(:,2)).*rmax]; % absolute saturation
    % summarize
    sample = [Lab,rgb,hsv_,SphLab,hueD,qSat];
    lgd = {'L*','a*','b','R','G','B','H','S','V','Hue (rad)','Elevation (rad)','Distance from Black','Reference distance from Black','Hue (deg)','abs. Sat.','abs. ref. Saturation (edge of gamut)'};
    % create table
    tbl = mat2cell(sample,ones(size(sample,1),1),ones(1,size(sample,2)));
    tbl = cell2table(tbl,"VariableNames",lgd);

end
%%
function [ft] = labFt(xXn)
    ft = NaN(size(xXn));
    for i = 1:numel(xXn)
        x = xXn(i);
        if x > ((6/29)^3)
            ft(i) = x^(1/3);
        else
            ft(i) = ((841/108)*x)+(4/29);
        end
    end
end
```
