## Supplementary Data S2 for "Identifying and engineering the molecular origins of lignin color for predictive staining of plant tissues": Human10DegStandardObserver.m

```
%% 10 degree spectral standard observer
stdOb = [360	0.0000001222	0.000000013398	0.000000535027
361	0.000000185138	0.000000020294	0.00000081072
362	0.00000027883	0.00000003056	0.0000012212
363	0.00000041747	0.00000004574	0.0000018287
364	0.00000062133	0.00000006805	0.0000027222
365	0.00000091927	0.00000010065	0.0000040283
366	0.00000135198	0.00000014798	0.0000059257
367	0.00000197654	0.00000021627	0.0000086651
368	0.0000028725	0.0000003142	0.000012596
369	0.0000041495	0.0000004537	0.000018201
370	0.0000059586	0.0000006511	0.0000261437
371	0.0000085056	0.0000009288	0.00003733
372	0.0000120686	0.0000013175	0.000052987
373	0.0000170226	0.0000018572	0.000074764
374	0.000023868	0.000002602	0.00010487
375	0.000033266	0.000003625	0.00014622
376	0.000046087	0.000005019	0.00020266
377	0.000063472	0.000006907	0.00027923
378	0.000086892	0.000009449	0.00038245
379	0.000118246	0.000012848	0.00052072
380	0.000159952	0.000017364	0.000704776
381	0.00021508	0.000023327	0.00094823
382	0.00028749	0.00003115	0.0012682
383	0.00038199	0.00004135	0.0016861
384	0.00050455	0.00005456	0.0022285
385	0.00066244	0.00007156	0.0029278
386	0.0008645	0.0000933	0.0038237
387	0.0011215	0.00012087	0.0049642
388	0.00144616	0.00015564	0.0064067
389	0.00185359	0.0001992	0.0082193
390	0.0023616	0.0002534	0.0104822
391	0.0029906	0.0003202	0.013289
392	0.0037645	0.0004024	0.016747
393	0.0047102	0.0005023	0.02098
394	0.0058581	0.0006232	0.026127
395	0.0072423	0.0007685	0.032344
396	0.0088996	0.0009417	0.039802
397	0.0108709	0.0011478	0.048691
398	0.0131989	0.0013903	0.05921
399	0.0159292	0.001674	0.071576
400	0.0191097	0.0020044	0.0860109
401	0.022788	0.002386	0.10274
402	0.027011	0.002822	0.122
403	0.031829	0.003319	0.14402
404	0.037278	0.00388	0.16899
405	0.0434	0.004509	0.19712
406	0.050223	0.005209	0.22857
407	0.057764	0.005985	0.26347
408	0.066038	0.006833	0.3019
409	0.075033	0.007757	0.34387
410	0.084736	0.008756	0.389366
411	0.095041	0.009816	0.43797
412	0.105836	0.010918	0.48922
413	0.117066	0.012058	0.5429
414	0.128682	0.013237	0.59881
415	0.140638	0.014456	0.65676
416	0.152893	0.015717	0.71658
417	0.165416	0.017025	0.77812
418	0.178191	0.018399	0.84131
419	0.191214	0.019848	0.90611
420	0.204492	0.021391	0.972542
421	0.21765	0.022992	1.0389
422	0.230267	0.024598	1.1031
423	0.242311	0.026213	1.1651
424	0.253793	0.027841	1.2249
425	0.264737	0.029497	1.2825
426	0.275195	0.031195	1.3382
427	0.285301	0.032927	1.3926
428	0.295143	0.034738	1.4461
429	0.304869	0.036654	1.4994
430	0.314679	0.038676	1.55348
431	0.324355	0.040792	1.6072
432	0.33357	0.042946	1.6589
433	0.342243	0.045114	1.7082
434	0.350312	0.047333	1.7548
435	0.357719	0.049602	1.7985
436	0.364482	0.051934	1.8392
437	0.370493	0.054337	1.8766
438	0.375727	0.056822	1.9105
439	0.380158	0.059399	1.9408
440	0.383734	0.062077	1.96728
441	0.386327	0.064737	1.9891
442	0.387858	0.067285	2.0057
443	0.388396	0.069764	2.0174
444	0.387978	0.072218	2.0244
445	0.386726	0.074704	2.0273
446	0.384696	0.077272	2.0264
447	0.382006	0.079979	2.0223
448	0.378709	0.082874	2.0153
449	0.374915	0.086	2.006
450	0.370702	0.089456	1.9948
451	0.366089	0.092947	1.9814
452	0.361045	0.096275	1.9653
453	0.355518	0.099535	1.9464
454	0.349486	0.102829	1.9248
455	0.342957	0.106256	1.9007
456	0.335893	0.109901	1.8741
457	0.328284	0.113835	1.8451
458	0.32015	0.118167	1.8139
459	0.311475	0.122932	1.7806
460	0.302273	0.128201	1.74537
461	0.292858	0.133457	1.7091
462	0.283502	0.138323	1.6723
463	0.274044	0.143042	1.6347
464	0.264263	0.147787	1.5956
465	0.254085	0.152761	1.5549
466	0.243392	0.158102	1.5122
467	0.232187	0.163941	1.4673
468	0.220488	0.170362	1.4199
469	0.208198	0.177425	1.37
470	0.195618	0.18519	1.31756
471	0.183034	0.193025	1.2624
472	0.170222	0.200313	1.205
473	0.157348	0.207156	1.1466
474	0.14465	0.213644	1.088
475	0.132349	0.21994	1.0302
476	0.120584	0.22617	0.97383
477	0.109456	0.232467	0.91943
478	0.099042	0.239025	0.86746
479	0.089388	0.245997	0.81828
480	0.080507	0.253589	0.772125
481	0.072034	0.261876	0.72829
482	0.06371	0.270643	0.68604
483	0.055694	0.279645	0.64553
484	0.048117	0.288694	0.60685
485	0.041072	0.297665	0.57006
486	0.034642	0.306469	0.53522
487	0.028896	0.315035	0.50234
488	0.023876	0.323335	0.4714
489	0.019628	0.331366	0.44239
490	0.016172	0.339133	0.415254
491	0.0133	0.34786	0.390024
492	0.010759	0.358326	0.366399
493	0.008542	0.370001	0.344015
494	0.006661	0.382464	0.322689
495	0.005132	0.395379	0.302356
496	0.003982	0.408482	0.283036
497	0.003239	0.421588	0.264816
498	0.002934	0.434619	0.247848
499	0.003114	0.447601	0.232318
500	0.003816	0.460777	0.218502
501	0.005095	0.47434	0.205851
502	0.006936	0.4882	0.193596
503	0.009299	0.50234	0.181736
504	0.012147	0.51674	0.170281
505	0.015444	0.53136	0.159249
506	0.019156	0.54619	0.148673
507	0.02325	0.56118	0.138609
508	0.02769	0.57629	0.129096
509	0.032444	0.5915	0.120215
510	0.037465	0.606741	0.112044
511	0.042956	0.62215	0.10471
512	0.049114	0.63783	0.098196
513	0.05592	0.65371	0.092361
514	0.063349	0.66968	0.087088
515	0.071358	0.68566	0.082248
516	0.079901	0.70155	0.077744
517	0.088909	0.71723	0.073456
518	0.098293	0.73257	0.069268
519	0.107949	0.74746	0.06506
520	0.117749	0.761757	0.060709
521	0.127839	0.77534	0.056457
522	0.13845	0.78822	0.052609
523	0.149516	0.80046	0.049122
524	0.161041	0.81214	0.045954
525	0.172953	0.82333	0.04305
526	0.185209	0.83412	0.040368
527	0.197755	0.8446	0.037839
528	0.210538	0.85487	0.035384
529	0.22346	0.86504	0.032949
530	0.236491	0.875211	0.030451
531	0.249633	0.88537	0.028029
532	0.262972	0.89537	0.025862
533	0.276515	0.90515	0.02392
534	0.290269	0.91465	0.022174
535	0.304213	0.92381	0.020584
536	0.318361	0.93255	0.019127
537	0.332705	0.94081	0.01774
538	0.347232	0.94852	0.016403
539	0.361926	0.9556	0.015064
540	0.376772	0.961988	0.013676
541	0.391683	0.96754	0.012308
542	0.406594	0.97223	0.011056
543	0.421539	0.97617	0.009915
544	0.436517	0.97946	0.008872
545	0.451584	0.9822	0.007918
546	0.466782	0.98452	0.00703
547	0.482147	0.98652	0.006223
548	0.497738	0.98832	0.005453
549	0.513606	0.99002	0.004714
550	0.529826	0.991761	0.003988
551	0.54644	0.99353	0.003289
552	0.563426	0.99523	0.002646
553	0.580726	0.99677	0.002063
554	0.59829	0.99809	0.001533
555	0.616053	0.99911	0.001091
556	0.633948	0.99977	0.000711
557	0.651901	1	0.000407
558	0.669824	0.99971	0.000184
559	0.687632	0.99885	0.000047
560	0.705224	0.99734	NaN
561	0.722773	0.99526	NaN
562	0.740483	0.99274	NaN
563	0.758273	0.98975	NaN
564	0.776083	0.9863	NaN
565	0.793832	0.98238	NaN
566	0.811436	0.97798	NaN
567	0.828822	0.97311	NaN
568	0.845879	0.96774	NaN
569	0.862525	0.96189	NaN
570	0.878655	0.955552	NaN
571	0.894208	0.948601	NaN
572	0.909206	0.940981	NaN
573	0.923672	0.932798	NaN
574	0.937638	0.924158	NaN
575	0.951162	0.915175	NaN
576	0.964283	0.905954	NaN
577	0.977068	0.896608	NaN
578	0.98959	0.887249	NaN
579	1.00191	0.877986	NaN
580	1.01416	0.868934	NaN
581	1.0265	0.860164	NaN
582	1.0388	0.851519	NaN
583	1.051	0.842963	NaN
584	1.0629	0.834393	NaN
585	1.0743	0.825623	NaN
586	1.0852	0.816764	NaN
587	1.0952	0.807544	NaN
588	1.1042	0.797947	NaN
589	1.112	0.787893	NaN
590	1.11852	0.777405	NaN
591	1.1238	0.76649	NaN
592	1.128	0.755309	NaN
593	1.1311	0.743845	NaN
594	1.1332	0.73219	NaN
595	1.1343	0.720353	NaN
596	1.1343	0.708281	NaN
597	1.1333	0.696055	NaN
598	1.1312	0.683621	NaN
599	1.1281	0.671048	NaN
600	1.12399	0.658341	NaN
601	1.1189	0.645545	NaN
602	1.1129	0.632718	NaN
603	1.1059	0.619815	NaN
604	1.098	0.606887	NaN
605	1.0891	0.593878	NaN
606	1.0792	0.580781	NaN
607	1.0684	0.567653	NaN
608	1.0567	0.55449	NaN
609	1.044	0.541228	NaN
610	1.03048	0.527963	NaN
611	1.016	0.514634	NaN
612	1.0008	0.501363	NaN
613	0.98479	0.488124	NaN
614	0.96808	0.474935	NaN
615	0.95074	0.461834	NaN
616	0.9328	0.448823	NaN
617	0.91434	0.435917	NaN
618	0.89539	0.423153	NaN
619	0.87603	0.410526	NaN
620	0.856297	0.398057	NaN
621	0.83635	0.385835	NaN
622	0.81629	0.373951	NaN
623	0.79605	0.362311	NaN
624	0.77561	0.350863	NaN
625	0.75493	0.339554	NaN
626	0.73399	0.328309	NaN
627	0.71278	0.317118	NaN
628	0.69129	0.305936	NaN
629	0.66952	0.294737	NaN
630	0.647467	0.283493	NaN
631	0.62511	0.272222	NaN
632	0.60252	0.26099	NaN
633	0.57989	0.249877	NaN
634	0.55737	0.238946	NaN
635	0.53511	0.228254	NaN
636	0.51324	0.217853	NaN
637	0.49186	0.20778	NaN
638	0.47108	0.198072	NaN
639	0.45096	0.188748	NaN
640	0.431567	0.179828	NaN
641	0.41287	0.171285	NaN
642	0.39475	0.163059	NaN
643	0.37721	0.155151	NaN
644	0.36019	0.147535	NaN
645	0.34369	0.140211	NaN
646	0.32769	0.13317	NaN
647	0.31217	0.1264	NaN
648	0.29711	0.119892	NaN
649	0.2825	0.11364	NaN
650	0.268329	0.107633	NaN
651	0.25459	0.10187	NaN
652	0.2413	0.096347	NaN
653	0.22848	0.091063	NaN
654	0.21614	0.08601	NaN
655	0.2043	0.081187	NaN
656	0.19295	0.076583	NaN
657	0.18211	0.072198	NaN
658	0.17177	0.068024	NaN
659	0.16192	0.064052	NaN
660	0.152568	0.060281	NaN
661	0.14367	0.056697	NaN
662	0.1352	0.053292	NaN
663	0.12713	0.050059	NaN
664	0.11948	0.046998	NaN
665	0.11221	0.044096	NaN
666	0.10531	0.041345	NaN
667	0.098786	0.0387507	NaN
668	0.09261	0.0362978	NaN
669	0.086773	0.0339832	NaN
670	0.0812606	0.0318004	NaN
671	0.076048	0.0297395	NaN
672	0.071114	0.0277918	NaN
673	0.066454	0.0259551	NaN
674	0.062062	0.0242263	NaN
675	0.05793	0.0226017	NaN
676	0.05405	0.0210779	NaN
677	0.050412	0.0196505	NaN
678	0.047006	0.0183153	NaN
679	0.043823	0.0170686	NaN
680	0.0408508	0.0159051	NaN
681	0.038072	0.0148183	NaN
682	0.035468	0.0138008	NaN
683	0.033031	0.0128495	NaN
684	0.030753	0.0119607	NaN
685	0.028623	0.0111303	NaN
686	0.026635	0.0103555	NaN
687	0.024781	0.0096332	NaN
688	0.023052	0.0089599	NaN
689	0.021441	0.0083324	NaN
690	0.0199413	0.0077488	NaN
691	0.018544	0.0072046	NaN
692	0.017241	0.0066975	NaN
693	0.016027	0.0062251	NaN
694	0.014896	0.005785	NaN
695	0.013842	0.0053751	NaN
696	0.012862	0.0049941	NaN
697	0.011949	0.0046392	NaN
698	0.0111	0.0043093	NaN
699	0.010311	0.0040028	NaN
700	0.00957688	0.00371774	NaN
701	0.008894	0.00345262	NaN
702	0.0082581	0.00320583	NaN
703	0.0076664	0.00297623	NaN
704	0.0071163	0.00276281	NaN
705	0.0066052	0.00256456	NaN
706	0.0061306	0.00238048	NaN
707	0.0056903	0.00220971	NaN
708	0.0052819	0.00205132	NaN
709	0.0049033	0.00190449	NaN
710	0.00455263	0.00176847	NaN
711	0.0042275	0.00164236	NaN
712	0.0039258	0.00152535	NaN
713	0.0036457	0.00141672	NaN
714	0.0033859	0.00131595	NaN
715	0.0031447	0.00122239	NaN
716	0.0029208	0.00113555	NaN
717	0.002713	0.00105494	NaN
718	0.0025202	0.00098014	NaN
719	0.0023411	0.00091066	NaN
720	0.00217496	0.00084619	NaN
721	0.0020206	0.00078629	NaN
722	0.0018773	0.00073068	NaN
723	0.0017441	0.00067899	NaN
724	0.0016205	0.00063101	NaN
725	0.0015057	0.00058644	NaN
726	0.0013992	0.00054511	NaN
727	0.0013004	0.00050672	NaN
728	0.0012087	0.00047111	NaN
729	0.0011236	0.00043805	NaN
730	0.00104476	0.00040741	NaN
731	0.00097156	0.000378962	NaN
732	0.0009036	0.000352543	NaN
733	0.00084048	0.000328001	NaN
734	0.00078187	0.000305208	NaN
735	0.00072745	0.000284041	NaN
736	0.0006769	0.000264375	NaN
737	0.00062996	0.000246109	NaN
738	0.00058637	0.000229143	NaN
739	0.00054587	0.000213376	NaN
740	0.000508258	0.00019873	NaN
741	0.0004733	0.000185115	NaN
742	0.0004408	0.000172454	NaN
743	0.00041058	0.000160678	NaN
744	0.00038249	0.00014973	NaN
745	0.00035638	0.00013955	NaN
746	0.00033211	0.000130086	NaN
747	0.00030955	0.00012129	NaN
748	0.00028858	0.000113106	NaN
749	0.00026909	0.000105501	NaN
750	0.000250969	0.000098428	NaN
751	0.00023413	0.000091853	NaN
752	0.00021847	0.000085738	NaN
753	0.00020391	0.000080048	NaN
754	0.00019035	0.000074751	NaN
755	0.00017773	0.000069819	NaN
756	0.00016597	0.000065222	NaN
757	0.00015502	0.000060939	NaN
758	0.0001448	0.000056942	NaN
759	0.00013528	0.000053217	NaN
760	0.00012639	0.000049737	NaN
761	0.0001181	0.000046491	NaN
762	0.00011037	0.000043464	NaN
763	0.00010315	0.000040635	NaN
764	0.000096427	0.000038	NaN
765	0.000090151	0.0000355405	NaN
766	0.000084294	0.0000332448	NaN
767	0.00007883	0.0000311006	NaN
768	0.000073729	0.000029099	NaN
769	0.000068969	0.0000272307	NaN
770	0.0000645258	0.000025486	NaN
771	0.000060376	0.0000238561	NaN
772	0.0000565	0.0000223332	NaN
773	0.00005288	0.0000209104	NaN
774	0.000049498	0.0000195808	NaN
775	0.000046339	0.0000183384	NaN
776	0.000043389	0.0000171777	NaN
777	0.000040634	0.0000160934	NaN
778	0.00003806	0.00001508	NaN
779	0.000035657	0.0000141336	NaN
780	0.0000334117	0.000013249	NaN
781	0.000031315	0.0000124226	NaN
782	0.000029355	0.0000116499	NaN
783	0.000027524	0.0000109277	NaN
784	0.000025811	0.0000102519	NaN
785	0.000024209	0.0000096196	NaN
786	0.000022711	0.0000090281	NaN
787	0.000021308	0.000008474	NaN
788	0.000019994	0.0000079548	NaN
789	0.000018764	0.0000074686	NaN
790	0.0000176115	0.0000070128	NaN
791	0.000016532	0.0000065858	NaN
792	0.000015521	0.0000061857	NaN
793	0.000014574	0.0000058107	NaN
794	0.000013686	0.000005459	NaN
795	0.000012855	0.0000051298	NaN
796	0.000012075	0.0000048206	NaN
797	0.000011345	0.0000045312	NaN
798	0.000010659	0.0000042591	NaN
799	0.000010017	0.0000040042	NaN
800	0.00000941363	0.00000376473	NaN
801	0.0000088479	0.00000353995	NaN
802	0.0000083171	0.00000332914	NaN
803	0.000007819	0.00000313115	NaN
804	0.0000073516	0.00000294529	NaN
805	0.000006913	0.00000277081	NaN
806	0.0000065015	0.00000260705	NaN
807	0.0000061153	0.00000245329	NaN
808	0.0000057529	0.00000230894	NaN
809	0.0000054127	0.00000217338	NaN
810	0.00000509347	0.00000204613	NaN
811	0.0000047938	0.00000192662	NaN
812	0.0000045125	0.0000018144	NaN
813	0.0000042483	0.00000170895	NaN
814	0.0000040002	0.00000160988	NaN
815	0.0000037671	0.00000151677	NaN
816	0.000003548	0.00000142921	NaN
817	0.0000033421	0.00000134686	NaN
818	0.0000031485	0.00000126945	NaN
819	0.0000029665	0.00000119662	NaN
820	0.00000279531	0.00000112809	NaN
821	0.0000026345	0.00000106368	NaN
822	0.0000024834	0.00000100313	NaN
823	0.0000023414	0.00000094622	NaN
824	0.0000022078	0.00000089263	NaN
825	0.000002082	0.00000084216	NaN
826	0.0000019636	0.00000079464	NaN
827	0.0000018519	0.00000074978	NaN
828	0.0000017465	0.00000070744	NaN
829	0.0000016471	0.00000066748	NaN
830	0.00000155314	0.0000006297	NaN];
```
