## Supplementary Data S2 for "Identifying and engineering the molecular origins of lignin color for predictive staining of plant tissues": readme.txt

This directory contains a script to compute CIE color spaces from UV-Vis-NIR spectra.
The workflow is contained in uvVisSpecColor.
This script was generated in MATLAB 2025b and may work with older versions of MATLAB.

Note: Necessary functions are located in the 'function' folder.
importCraicSpecfile … initial import of CSV data from the microscope used in our publication.
importUVvis… transforms raw import into UV-Vis spectrum for multiple light sources and combines them.
spec2cie … converts transmission data into various color spaces, like L*a*b*, rgb, hsv, etc.

Note: This script is customised to work with input data generated from exports of CRAIC microscopy software and may not import other CSV files correctly. The main function 'spec2cie' works with any variable in MATLAB as long as the workspace has a variable that has transmission spectra as m x 1 vectors and a compatible wavelength vector with m wavelengths of the same dimensions.
