## Supplementary Data S2 for "Identifying and engineering the molecular origins of lignin color for predictive staining of plant tissues": uvVisSpecColor.m

```
%% get Color data from spectrum
close all
clear

addpath('.\function');
%% Define directory for your data when loading exported CSV files from 
% CRAIC software.
% The directory should only contain csv files of UV-Vis spectra.

path = 'C:\ your directory'; % ADD your folder here!!!
%%% or
path = fullfile('test');

% example: path =
% 'E:\CRAIC\TissueSections\pointMeas\wt_20251118\glassslides';
%% load CRAIC csv files
ds = datastore(path,'Type','tabulartext','FileExtensions','.csv');
ds = ds.Files;
n = numel(ds);
%%% create library variable
specLib = cell(numel(ds),4);
for i = 1:n
    %%%% output spectra [sum(UV,Vis], UV, Vis]
    [absor,trans,wavelength,lamp,~] = importUVvis(ds{i});
    %%%% clean up inf values
    for j = 1:3
        isInf = isinf(absor(:,j));
        absor(:,j) = interp1(wavelength(~isInf),absor(~isInf,j),wavelength);
        isInf = isinf(trans(:,j));
        trans(:,j) = interp1(wavelength(~isInf),trans(~isInf,j),wavelength);
    end
    specLib(i,:) = {absor, trans, wavelength,lamp};
end
clearvars -except specLib ds path n wavelength
%% get Color data
display = true; % generate one figure per sample spectrum. Set to 'false' if not wanted.

ls = 1; % 1 ... sum of UV and Vis lamp, 2 ... UV lamp only, 3... Vis lamp only
useE = true;    % set to 'true' to use ideal light source, 
% 'false' will use spectrum specific light source
lamp = ones(numel(wavelength,1));

%%% settings for spectrum type
norm = true; % set to 'false' to get raw spectrum colors
range = [280,1200]; % set spectral range in which to normalize spectral data
normType = 1;       % 1... range normalization (default), 2... standard normalization

normLgd = {'raw spectrum','range norm.', 'standard norm.'};

%%% output matrices
specLab = NaN(n,3);     % L*a*b* matrix
specHueD = NaN(n,1);    % hue angle matrix
specQSat = NaN(n,2);    % quantitative chroma/saturation matrix, 2nd value is gamut edge
specSphLab = NaN(n,4);  % custom spherical representation of L*a*b* as (hue(radians), elevation(radians) from black, distance from black]
specRGB = NaN(n,3);     % rgb code matrix
specHSV = NaN(n,3);     % hsv code matrix
specEdit = specLib;     % used modified spectrum
specEdit(:,4) = {'raw spectrum'};

%%% get data
for i = 1:n
    if ~useE
        lamp = specLib{i,4}(:,ls);
    end
    transRel = specLib{i,2}(:,ls);
    wavelength = specLib{i,3}(:,ls);

    if norm
        idxRange = wavelength>range(1) & wavelength<range(2);
        minT = quantile(transRel(idxRange,:),0.001);
        maxT = quantile(transRel(idxRange,:),0.999);
        if normType == 2        % standard normalization
            transRel = (transRel - minT)./std(transRel);
        else                    % range normalization
            transRel = (transRel - minT)./(maxT-minT);
            transRel = transRel.*0.998 + 0.001;
        end
        specEdit(i,4) = normLgd(:,normType + 1);
        specEdit{i,2} = transRel;
        
    end
    absor = real(-log10(transRel));
    specEdit{i,3} = absor;

    [Lab,hueD,qSat,SphLab,rgb,hsv_,tbl] = spec2cie(transRel,wavelength,lamp);
    %%% OPTIONAL %%%%%%%%%%%%%%%%%%%%%%%%%%%%%%%%%%%%%%%%%%%%%%%%%%%%%%%%%%
    if display
        f = figure(i);
        theme(f,'light');
        s = subplot(2,1,1);
        plot(wavelength,transRel.*100);
        ylabel({specEdit{i,4};'T(%)'})
        xlabel('lambda [nm]');
        xBlock = 0.05*diff(range);
       
        xlim(range + [0,xBlock]);
        try
            hold on;
            dispClr = [lab2rgb(Lab,"WhitePoint","e");...
                hsv2rgb([hsv_(1),1,1]);...
                [1,1,1]-hsv_(2);...
                [1,1,1].*hsv_(3)];
            dispClr(dispClr<0) = 0;
            dispClr(dispClr>1) = 1;
            dispClr = permute(dispClr,[1,3,2]);
            imagesc(range(2)+[xBlock*0.5,xBlock],[25,100]-12.5,dispClr);
            hold off;
            text(range(2)+ones(4,1),[25;50;75;100]-12.5,{'RGB  ','H  ','S  ','V  '},'HorizontalAlignment','right');
        catch
        end

        s2 = subplot(2,1,2);
        plot(wavelength,absor);
        ylabel({specEdit{i,4};'Abs. [a.u.]'})
        xlabel('lambda [nm]');
        xlim(range);
        ylim([0,2]);
    end
    %%%%%%%%%%%%%%%%%%%%%%%%%%%%%%%%%%%%%%%%%%%%%%%%%%%%%%%%%%%%%%%%%%%%%%%
    if i == 1
        colorTbl = tbl;
    else
       colorTbl = [colorTbl;tbl];
    end
end
modTbl = table(categorical(specEdit(:,4)),'VariableNames',{'Type'});
filesTbl = table(ds,'VariableNames',{'File path'});
colorTbl = [filesTbl,modTbl,colorTbl];

clearvars -except color** specLib ds path n wavelength spec**
   
%% export table
writetable(colorTbl,fullfile(path,'colorData.xls'),'Sheet','ColorAnalysis','Range','A2');
```
